# The “Proxitome” of Arabidopsis Processing Bodies Reveals a Condensate-Membrane Interface Instructing Actin-Driven Directional Growth

**DOI:** 10.1101/2022.06.03.494746

**Authors:** Chen Liu, Andriani Mentzelopoulou, Amna Muhammad, Andriy Volkov, Dolf Weijers, Emilio Gutierrez-Beltran, Panagiotis N. Moschou

## Abstract

Cellular condensates can comprise membrane-less ribonucleoprotein assemblies with liquid-like properties. These cellular condensates influence various biological outcomes, but their liquidity hampers their isolation and characterization. Here, we investigated the composition of the condensates known as processing bodies (PBs) in the model plant Arabidopsis through a proximity-biotinylation pipeline. Using *in situ* protein–protein interaction approaches, genetics and high-resolution imaging, we show that PBs comprise networks that interface with membranes. Surprisingly, the conserved component of PBs, DECAPPING PROTEIN 1 (DCP1), can interface with unique plasma membrane subdomains that include cell edges and vertices. We characterized these plasma membrane interfaces and discovered a developmental module that can control cell shape. This module is modulated by the liquid-like properties of DCP1 and the actin-nucleating SCAR–WAVE complex, whereby the DCP1-SCAR–WAVE interaction confines actin nucleation. This study reveals an unexpected repertoire for a conserved condensate at unique membrane-condensate interfaces.

## Introduction

Proteins may participate in multivalent interactions with themselves or one another. Multivalency can depend on weak interactions between charged residues, dipoles, and/or aromatic groups often displayed by so-called “intrinsically disordered regions” (IDRs). These interactions can promote their dissociation from the bulk protein pool to form droplet-like condensates through, for example, liquid–liquid phase separation (LLPS)^1^. As such, LLPS is a state favoring condensation through weak intra- or intermolecular interactions. We use the term “condensates” hereafter to describe such proteinaceous assemblies. LLPS occurs when the bulk concentration exceeds a threshold above which molecules spontaneously partition into droplets. Transient or more stable condensates form in the nucleus, cytoplasm or plasma membrane (PM) interfaces, e.g., the animal-specific junction adherent molecules^2, 3^.

Condensates can coarsen, increase or decrease in size over time on membrane surfaces^4^. Likewise, the archetypal and evolutionarily conserved condensates known as processing bodies (PBs), which are rich in proteins and RNA and modulate RNA silencing, decapping and decay, undergo fission at membrane surfaces of the endoplasmic reticulum (ER) through an unknown mechanism^5^. Although systematic studies of plants PBs are lacking, animal PBs appear to be long-term storage sites for mRNAs poised to be released for translation following specific cues related to stress, metabolism and translation capacity.

Albeit readily visible in cells, as aforemenioned, condensates such as PBs depend on a meshwork of weak interactions and lack a surrounding membrane, which makes their isolation challenging. Proximity-dependent biotin ligation (or PDL) harnesses covalent biotinylation of interacting proteins with or near neighbors of a particular prey protein; we and others have recently established PDL approaches in various plants^6–14^. By bypassing the need to retain native interactions for their identification, PDL holds promise to delineate the organizational and functional principles of condensates.

Inspired by PDL applications for the elucidation of condensates in non-plant models^15^, we established a PDL pipeline to determine the protein composition of condensates in plants, using PBs as a proof of concept. We show here that PBs are organized into both known and unknown functional modules. We further discovered that PBs interface with plasma membrane (PM) domains and cell edges or vertices. The suppressor of the cAMP receptor (SCAR)-WASP family verprolin homologous (WAVE) complex modulates this interface, which results in actin nucleation and a coordinated cellular system that can culminate in growth regulation.

## Results

### A bait for PB proteome capture

To determine the composition of condensates in plants, we focused on PBs as a proof of concept, as non-plant PBs are omnipresent and share some features with LLPS condensates. Depending on the conditions, PBs show dynamic behavior and grow, shrink (through fission) or fuse like membrane-bound organelles^16^. We dynamically imaged the cellular behavior of the evolutionarily conserved PB component DECAPPING PROTEIN 1 (DCP1) fused to green fluorescent protein (GFP; Supplementary Fig. 1a, b and Supplementary File 1) by confocal microscopy. We placed the expression of the *DCP1-GFP* construct under the control of the *DCP1* promoter or two constitutive promoters. We confirmed that plant PBs exhibit the same features as their animal counterparts (Supplementary Figs. 1b,c and 2 and ref.17). Indeed, GFP-positive puncta showed fusion, fission and alterations in size or shape with high circularity, suggesting that: 1) they behave like liquid droplets with high surface tension, and 2) they form by LLPS. Notably, when expressed under the control of the strong and constitutive cauliflower mosaic virus (CaMV) *35S* promoter, DCP1-positive puncta exhibited more variable circularity (Supplementary Figs. 1c, 2c) and reduced fusion and fission (Supplementary Fig. 1c) compared to those produced with two other promoters (*DCP1pro* or *RPS5apro* from *RIBOSOMAL PROTEIN S5a*). This suggested that elevated DCP1 concentration due to overexpression leads to a coarsening of PBs. Together, these results confirmed that plant PBs have features of LLPS condensates and can coarsen depending on the saturation concentrations of their components.

We next established a stepwise PDL-based approach. We postulated that incorporating a standard affinity purification (AP) step, using anti-FLAG beads to capture a FLAG-tagged protein, before a subsequent PDL step (with the biotin ligase “TurboID”; pipeline in Fig. 1a and detailed protocol in Methods) might help identify strong or direct (AP) versus more transient and weak or indirect interactions (PDL; hereafter defined as the “proxitome”) with DCP1. To this end, we used a construct encoding a chimeric DCP1-TurboID-6x**<u>H</u>**is-3x**<u>F</u>**LAG (*DCP1-TurboID-HF*), with superfolder-GFP as a control, whose encoding sequence was driven by the *35S* promoter (“noise”; *GFP-TurboID-HF*). DCP1-TurboID-HF was functional, showing higher expression levels (∼20%) than were seen in wild-type (WT) seedlings, and could rescue the adult phenotype of the weak *dcp1-3* mutant^18^ (Supplementary Fig. 3a-c; see also below). These data suggested that DCP1-TurboID-HF is a physiologically relevant bait with which to explore the PB interactome.

**Fig. 1.**
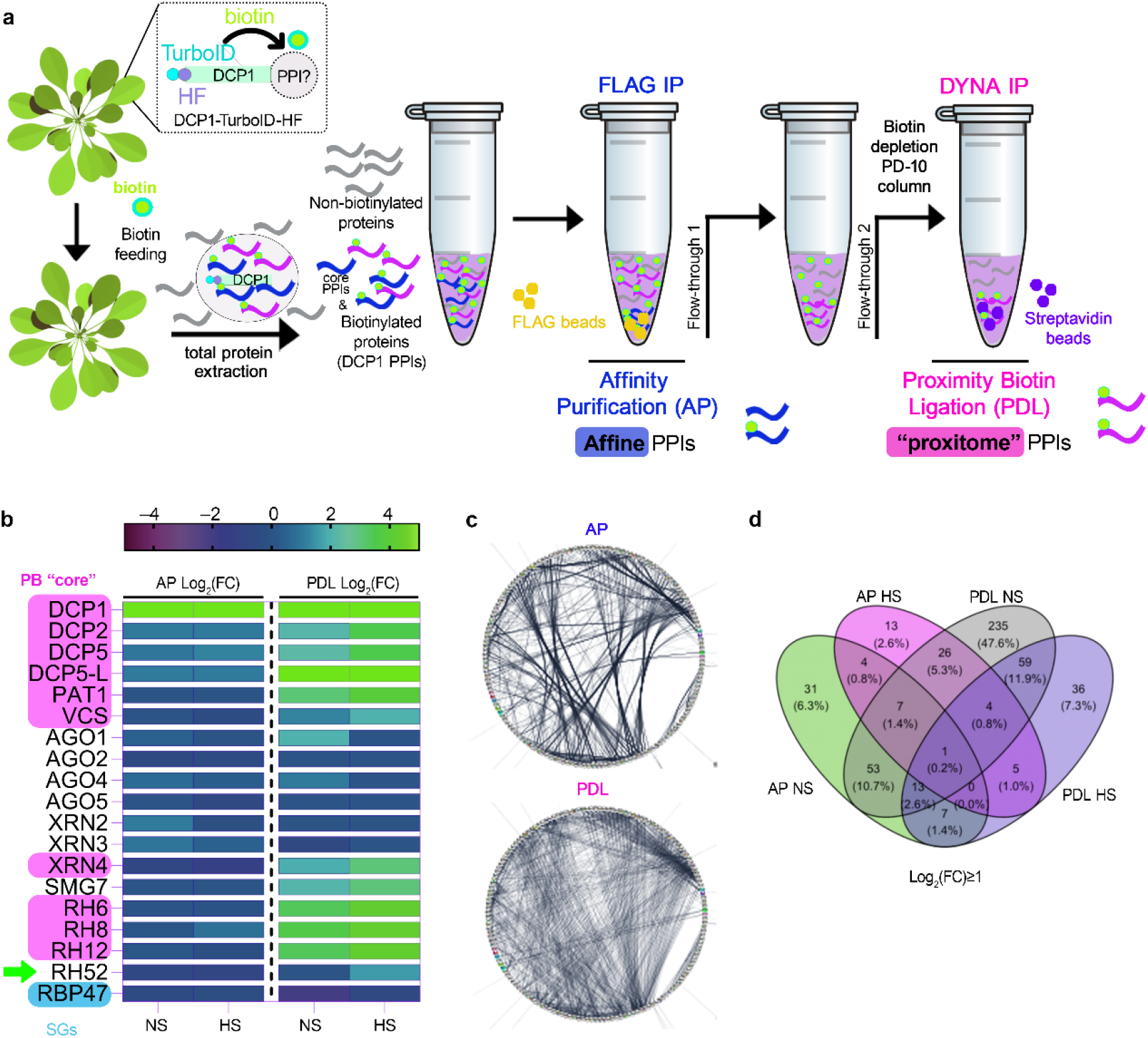
The pipeline of the APEAL approach. **a,** Overview of the APEAL pipeline. Upon 24 h biotin feeding and treatment (supply 50 μM biotin directly into leaves of 4-week-old plants by syringe infiltration), total proteins are extracted from infiltrated leaves. The proteome is subjected to AP-immunocapture of the FLAG tag. In the AP step, some of the captured proteins will be biotinylated. The PDL step uses the leftover supernatant from the AP step and captures biotinylated proteins with streptavidin beads. PPI, protein protein interactions; AP, affinity purification; PDL, proximity-dependent biotin ligation; FLAG IP, FLAG-beads immunoprecipitation; DYNA-IP, streptavidin-bead immunoprecipitation. We use the term “proxitome”, to describe the proteins captured by the PDL step of APEAL. These proteins may not necessarily physically interact with DCP1. **b,** Heatmap showing the “core processing body (PB)” (magenta) components identified and other linked proteins. RBP47 is a stress granule marker (SGs; blue). RH52 is a new helicase identified as a PB component (green arrow). The scale on the right shows log_2_FC of protein abundance. Note that only the PB core components were enriched in the PDL (i.e., log_2_FC∼1 or above). Furthermore, heat stress (HS) increased the enrichment of PB core components. **c,** Comparison of AP/PDL interacting networks produced from APEAL. STRING density plots of pairwise interactions between proteins obtained from the AP or PDL steps (combined interactions found in non-stress [NS]/[HS]). Note that PDL produces an overall denser interaction network (under standard parameters, the same number of proteins was selected for AP/PDL; see main text also). **d** Venn diagram showing the proteins identified for PDL and AP in NS and HS samples (PPIs fulfilling the criterion log_2_FC>1).

### APEAL captured the PB proteome

We hoped that the AP/PDL combination might capture both direct and stable interactions but also the proxitome, which could be more relevant for condensates, as they form through rather weak interactions. After the AP step, we detected biotinylated proteins in the remaining supernatant (defined as “flow-through 1” in Fig. 1a, Supplementary Fig. 4). We thus further analyzed proteins obtained from both the AP and PDL steps; this tandem analytical approach is referred to hereafter as “APEAL.” We coupled APEAL with nano-liquid chromatography tandem mass spectrometry (nano-LC-MS/MS) in biological duplicates on DCP1 baits using 4-week-old rosettes grown under heat stress (HS) conditions (37°C for 2 h) and non-stress (NS) conditions. We selected HS treatment to benchmark APEAL because PBs fuse and coarsen during HS (Supplementary Fig. 1c), which may lead to increases in their size and number and likely to a change in composition^5^. HS also enhances RNA decapping through the increased association of the mRNA decapping complex DCP1–DCP2 with polysomes^19^. We observed that HS does not significantly affect biotinylation levels of DCP1-TurboID-HF (Supplementary Fig. 3b; compare “24 h” vs. “HS”), suggesting that any differences in the PB proteome between NS and HS conditions may be of biological relevance.

Proteome analyses yielded almost equivalent numbers of protein hits for AP and PDL steps (Supplementary Fig. 5; without filtering). Unexpectedly, and despite the size increase of PBs noted above, HS led to fewer proteins being captured in PDL/HS from DCP1-TurboID-HF, suggesting that DCP1 may be spatially more confined, perhaps within PBs (see below for a possible explanation), thus limiting the capture of its proxitome. To test for enrichments across the different samples, we applied the same criteria described in our previous work to define high-confidence proximity interactions^6^. Briefly, we considered only high-confidence interactions present in both biological replicates (log_2_[fold change]≥1, *n*=2, false-discovery rate [FDR]=0.05; Supplementary File 2). To compare the AP and PDL datasets, we assigned LFQ (label-free quantitation, Supplementary File 2a) and iBAQ values (intensity-based absolute quantification, obtained by dividing protein intensities by the number of theoretically observable tryptic peptides between 6 and 30 residues; Supplementary File 2b) to the semi-quantitative peptide enrichments and reiterated interactions. The LFQ/iBAQ assignments produced similar results (Supplementary File 3).

An inspection of protein hits in the PDL step confirmed an enrichment for conserved PB core components (Fig. 1b; log_2_FC˃1)^19^. As a cautionary note, we do not define the PB core as a structural entity with topological essence (e.g., with the PB center formed as an immiscible liquid), but rather as an indispensable set of proteins that comprises part of the decapping machinery. Intriguingly, the AP step failed to enrich for PB core components, suggesting that many weak interactions are not retained or protein subcomplexes aggregate (e.g., condensate) and precipitate during the immunopurification steps, and thus validating our dual approach (Fig. 1b; “AP” vs. “PDL”). Importantly, APEAL showed no enrichment for proteins exclusively associated with the other condensates known as stress granules (SGs; e.g., RNA-BINDING PROTEIN 47B [RBP47]), despite the physical links between the two condensates^19^. One exception was the class of RNA helicases present in both SGs and PBs (Fig. 1b; log_2_FC≥1.5; Fig. 1b, RH6, RH8, RH12 and the newly discovered RH52 herein). The absence of SG components confirmed that APEAL can function at a submicron scale, at least in this case, thereby providing the required spatial resolution to specifically resolve the composition of condensates.

### Network analyses reveal that PBs interface with membranes

We integrated our PDL hits with delimited direct protein–protein interaction (PPI) data from the STRING database and generated PPI networks, revealing that the PDL dataset forms a denser PPI network than that obtained with AP. The average numbers of interactions from PDL per protein were 5.4 (*p*=5.6e^−11^) and 4.1 (*p*=2.1e^−10^) for NS and HS, respectively (Fig. 1c). The PDL dataset also showed a different profile in terms of interactions from AP, while different interactions were observed when HS and NS were compared (Fig. 1d). Consistent with the expected incremental formation of PBs during HS, the term “PΒs” was more enriched in the PDL dataset upon HS when compared to NS (*p*=6.9e^−2^ vs. 6.9e^−22^). We detected a partial overlap between our datasets and other published proteomics datasets obtained by AP: PM (overlap 27.1%; datasets in Supplementary File 4), endomembranes (e.g., multivesicular bodies [MVBs], clathrin; overlap 9–32.8%) and DCP1–DCP2 (overlap of up to 36.3%)^20^. Notably, the results from the AP of DCP1 and DCP2 showed a lower-than-expected overlap with our datasets and a lack of membrane-related proteins. Although in the previous DCP1-DCP2 AP study formaldehyde was used, because it has the shortest span (∼2–3 Å) of any known crosslinker^21^, it may not be the best option to capture condensates that can potentially interface with membranes. More crosslinkers with various spans will need to be tested, but we speculate for now that PDL may expand the proteome coverage, as it allows the use of harsher conditions during purification (as we confirmed below). Furthermore, our datasets suggested that some PB components (but not the PB entity as such; see below) may interface with membranes, as the overlaps with the PM and the endomembrane datasets were comparable to those for the DCP1–DCP2 datasets; we clarify this in detail below. We further confirmed that the PDL step could predict interactions between newly identified components and DCP1 (Supplementary Fig. 6a,b; 9 out of 9 proteins tested).

We tested hierarchical linkages between the networks we generated by integrating the data from STRING with our datasets using hypergeometric tests (FDR=0.05). We successfully assigned four dense and interconnected subnetworks (Supplementary Fig. 2a-f) to PBs according to overrepresentations in AP and PDL datasets from both HS and NS conditions (Supplementary Fig. 7). Subnetwork 1 is related to RNA metabolism. In subnetwork 2, we obtained hits for phytohormone signaling, defense attenuation, translation and metabolism. Subnetwork 3 comprises an unexpected network for PBs, with hits linked to membrane remodeling and trafficking; finally, subnetwork 4 includes another surprising network of proteins linked to actin remodeling (Fig. 2f; Supplementary Files 5, 7). In Supplementary Fig. 7, we succinctly describe these networks, from which testable hypotheses may arise for future studies.

**Fig. 2.**
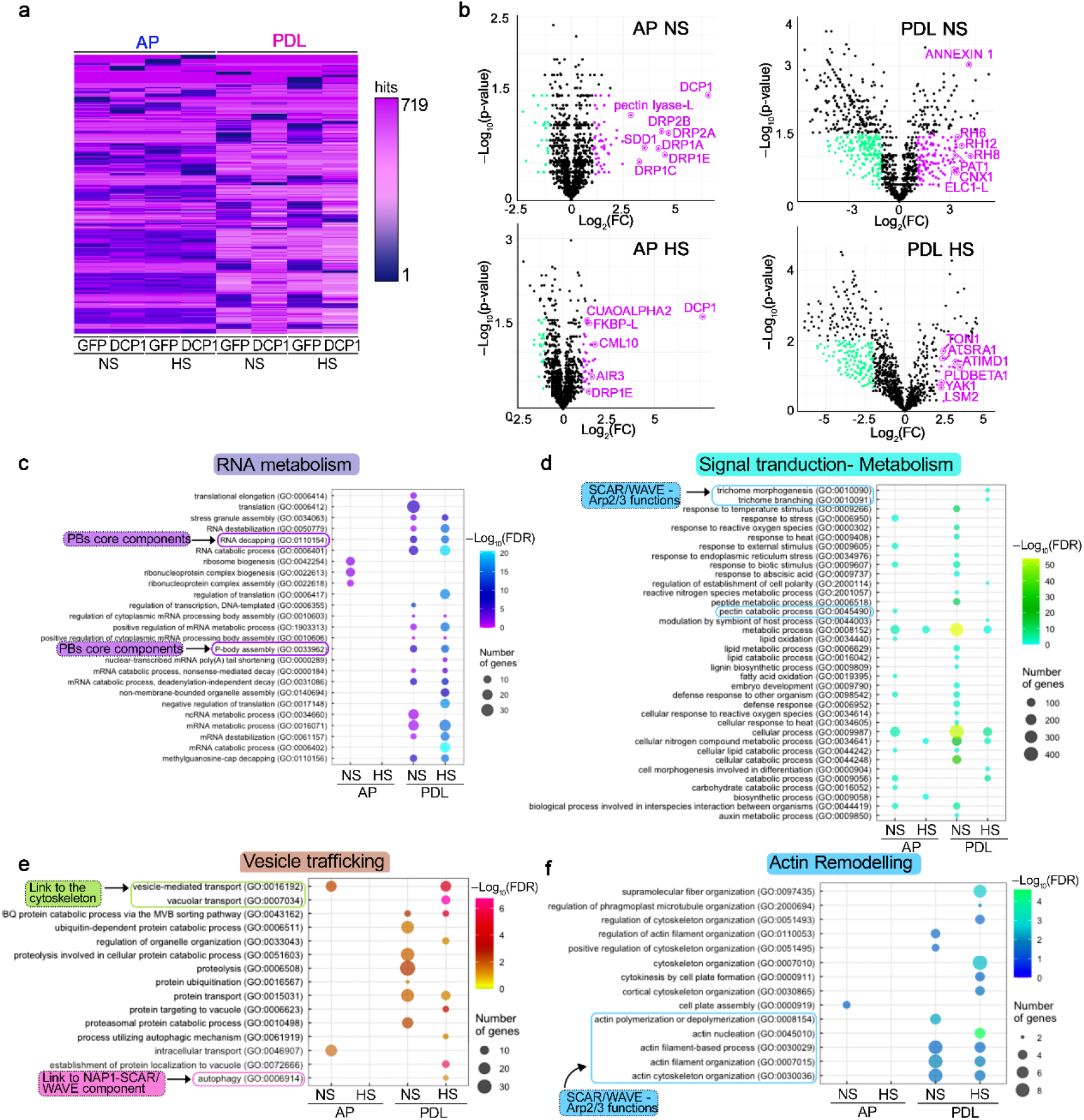
APEAL efficiently purifies PB components. **a,** Heatmap showing the different peptide enrichment in AP and PDL APEAL steps under NS and HS conditions. Note the different clusters in the different samples. The scale shows peptide number. **b,** Volcano plots showing significantly enriched proteins in NS and HS conditions from the AP and PDL APEAL steps. Selected proteins are indicated in magenta and are encoded by genes that belong to the identified subnetworks described in c,d. Magenta indicates enrichment in HS samples; cyan indicates depletion in HS samples. **c-f,** Gene Ontology (GO) enrichment analyses of the APEAL results, divided into four subnetworks. Note the terms related to vesicle trafficking and actin remodeling. A more detailed description is provided in Supplementary Fig. 7. Note that signal transduction proteins, metabolism-related proteins and vesicle trafficking proteins evade PBs during HS (Supplementary Fig. 5, reduced hit number), while the opposite pattern is observed for actin and RNA metabolism subnetworks. FDR, false discovery rate. Important links described below are indicated (i.e., to the SCAR–WAVE/Arp2–Arp3 (and the component NAP1), cytoskeleton, PB core and cell-wall-related metabolism; see also below for more details and Supplementary Fig. 7).

### DCP1 interfaces with plasma membrane domains at the cellular face independently of its role in decapping

We asked whether the links between PBs and subnetworks 3 and 4, which suggested endomembrane and cytoskeleton functions, respectively, might be physiologically relevant. We thus examined potential DCP1 connections with the endomembrane system and cytoskeleton. Total internal reflection microscopy (TIRFM), which is suitable for viewing cell surface processes due to shallow illumination penetration (decay constant ∼100 nm), allowed us to observe occasional stalling and fusion of DCP1-GFP-positive droplets with one another at the PM interface of meristematic root epidermal cells (Fig. 3a,b). Though encouraging, these results do not strongly support a direct association between DCP1 and PM lipids. As a cautionary note also, the lateral membranes used for imaging may not reflect the exact situation of apicobasal membranes during droplet dynamics. In addition, mean squared displacement (MSD) analyses, which track particles in two dimensions, showed that DCP1 droplets become more stable with time, after an initial Brownian-motion diffusion at the PM, and could also undergo coarsening (Fig. 3b, inset; track analyses). This result suggested that DCP1 droplets may coalesce to form larger 2D domains (i.e., at the inner face of the PM). Also of note is that PBs are more dynamic at the PM plane than in the cytoplasm (i.e., Fig. 3b vs. Supplementary Fig. 1b; “stable”), suggesting that the PM promotes PBs dynamics, manifested as fission (mainly in this case) or fusion.

**Fig. 3.**
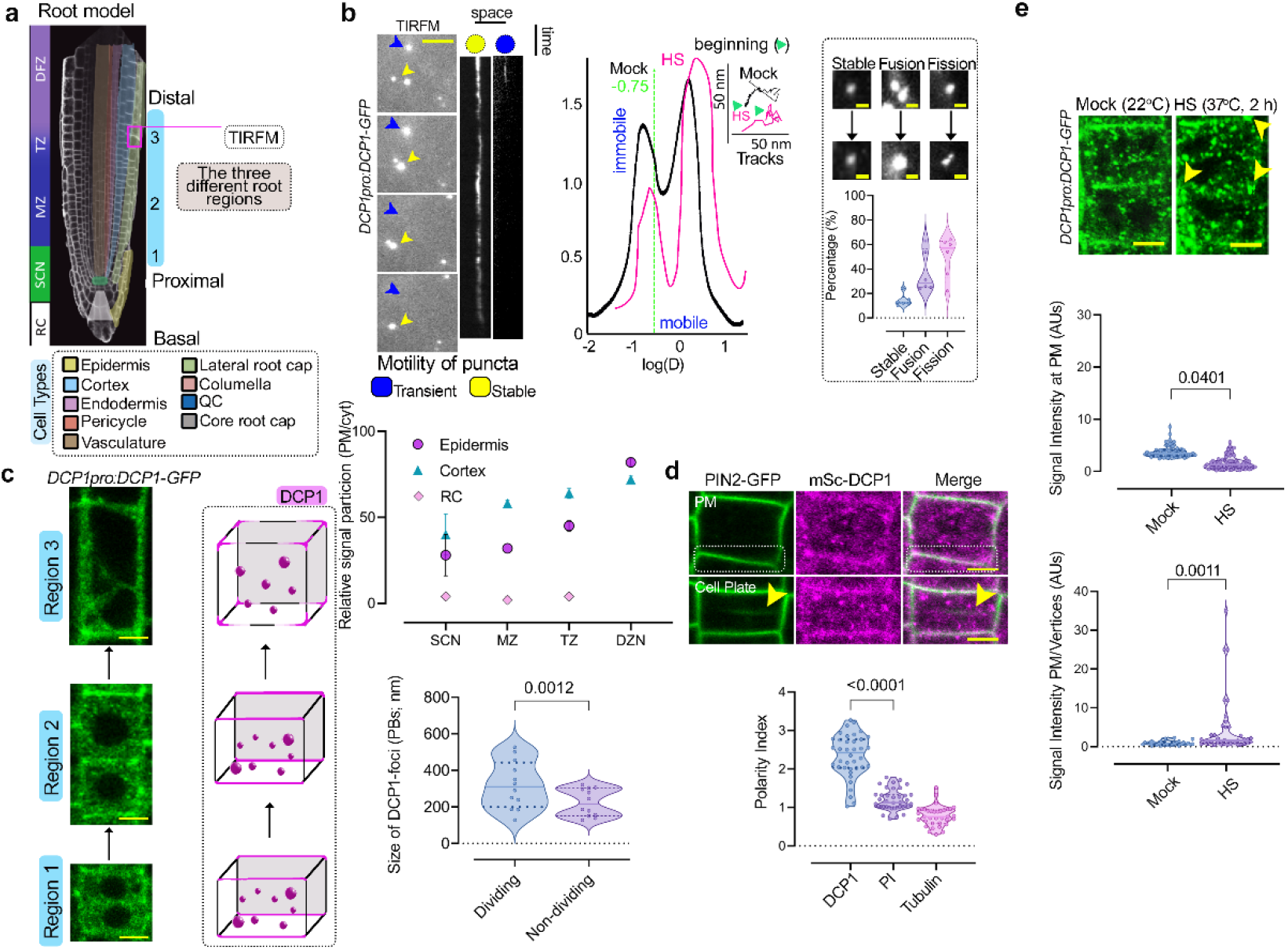
DCP1 protein interfaces with the plasma membrane. **a** Diagram of a root showing the three developmental regions (1–3) under examination in this study: stem cell niche (SCN; region 1), meristematic zone (MZ; region 2) and transition zone (TZ)-differentiation zone (DFZ; region 3). The different cell types are color-coded. The region used in TIRF-M experiments is highlighted by the dashed magenta rectangle (see also B). **b** Representative TIRF-M of DCP1-GFP (*DCP1pro:DCP1-GFP* transgene) in the lateral PM of root epidermal cells showing the transient attachment of DCP1-positive puncta to the PM. Yellow arrowheads denote a PB that shows motility at the PM focal plane; blue arrowheads show a PB that transiently localizes to the PM. The corresponding kymographs are shown to the right. Scale bars, 2 μm (space); 3 sec (time) for the kymograph. Middle: distribution of immobile and mobile DCP1 molecules relative to the motility Log(D) value of –0.75 (threshold), in NS or HS conditions (D, diffusion coefficient). Inset: individual trajectories of mobile DCP1 molecules in NS and HS conditions (500 frames, *n*=120), showing a combination of directional and Brownian motion for both NS/HS. The yellow arrowhead denotes the beginning of the track. Right: TIRF-M images showing stable DCP1-positive puncta or puncta undergoing fusion/fission, and frequency of non-dynamic (stable) or fusion/fission events (*N*=3, *n*=8–10). **c** DCP1 localization is developmentally regulated. Gradual edge or vertex accumulation of DCP1-GFP in three different root regions (left). Scale bars, 7 μm. Right: an average DCP1 behavioral model for localization in each region. Note that edge/vertex localization becomes more significant in region 3 (see below for robustness test). Bottom: relative signal intensity partitioning of DCP1 (PM/cyt; *N*=6, *n*=10–11) between the different root regions as described in (a) and PB size between dividing and non-dividing root cells (*N*=1, *n*=12, significant difference determined by Wilcoxon test). RC, root cap. **d** DCP1 is polarly localized. Representative confocal micrographs from lines co-expressing *RPS5apro:HF-mScarlet-DCP1* and *PIN2pro:GFP-PIN2* (epidermal cells, region 3). Scale bars, 3 μm. Bottom: polarity index of DCP1 in root meristematic cells (compared to propidium iodide and tubulin staining of root cells, not shown). Significant differences were determined by the Brown-Forsythe and Welch-ANOVA (*N*=3, *n*=20 cells). **e** Representative confocal micrograph of DCP1-GFP in root meristematic cells under NS/HS conditions (region 2). Scale bars, 5 μm. Note the depletion of DCP1 from the PM upon HS, but the increased edge/vertex signal.

High-speed super-resolution confocal microscopy (∼120 nm axial resolution) of micrographs and fluorescence partition analyses between the PM and the cytoplasm (overall signal) detected a fraction of DCP1-GFP at the PM, regardless of the promoter driving the encoding construct; this localization gradually increased along the root proximodistal axis (Fig. 3c). Notably, the ratio between the PM and cytosolic fluorescence signals indicated that cells with sharp edges accumulate more overall DCP1-GFP at cell edges (Fig. 3c). We use the term “edge” in the geometric sense of an intersection between two faces of a polyhedron, rather than of a periphery or front^22^. In dividing cells, PBs were globally more and larger (Fig. 3c). Other core PB components were associated with the PM either very transiently (in dividing cells) or not at all (e.g., DCP2; Supplementary Fig. 8a). None of these proteins accumulated at the cell edges. We further observed that DCP1 exhibits a relative polar PM localization, exemplified by the polar PIN-FORMED2 (PIN2) transporter, in the apicobasal domains (Fig. 3d). Under HS conditions, DCP1-GFP abundance decreased slightly at the PM in epidermal meristematic cells, but increased along the cell edges (Fig. 3e). Moreover, addition of cycloheximide (CHX, known to dissolve PBs^5^) decreased PB numbers and led to greater DCP1 localization at the PM (Supplementary Fig. 8b). Taken together, these results indicate that DCP1 localization at the PM might be reversible, while the dissolution of PBs followed by the rapid redirection of DCP1 to the PM suggested competition between PBs and membranes for DCP1. The transient association of some PB components was consistent with a model in which PBs undergo fission at the PM interface and only few components, such as DCP1, persist there.

Given the rapid and conditional DCP1 dissociation from the PM, which may also explain the decrease of proteins in APEAL relevant to subnetworks 3 and 4 (upon HS), we postulated that DCP1 localizes close to the cytoplasmic PM face and might not necessarily be inserted in the PM. We first tested whether DCP1 accumulates at the PM through the secretory pathway by incubating seedlings with 50 µM brefeldin A (BFA), a fungal toxin that prevents vesicle formation for exocytosis by inhibiting ADP ribosylation factor GEFs (ARF-GEFs). DCP1 did not accumulate intracellularly upon BFA treatment (Supplementary Fig. 8c), and it lacks a signal peptide or a transmembrane domain, suggesting that it is not inserted in the PM. These results suggested that DCP1 may use another route to reach the cytoplasmic PM face.

Dynamic 3D imaging of DCP1-GFP along the developmental root axis showed that DCP1 accumulates more extensively at cell edges later in development, when it is further restricted at vertices (Fig. 4a). Hence, this effect was more evident far from the quiescent center (QC) and was independent of the presence of PBs, as this accumulation was unaffected by the presence of CHX (Supplementary Fig. 8c); many cells displayed a single edge decorated with DCP1. Later in development, mainly at the transition zone, the DCP1 localization domain became further restricted, mainly decorating vertices (Fig. 4a; circular plots). The term “vertex” here is used in the geometric sense to define the internal angular point of a polygon and defined as an intersection between the three cell edges (cytoplasmic PM face). We observed DCP1 localization largely at the vertex in epidermal cells, a pattern unique to DCP1 among the core PB components tested (Supplementary Fig. 8a). Notably, this localization was reminiscent of that of mammalian LLPS zonula occludens (ZO) proteins, which form LLPS condensates in animal tricellular junctions, where they modulate epithelial integrity^23^.

**Fig. 4.**
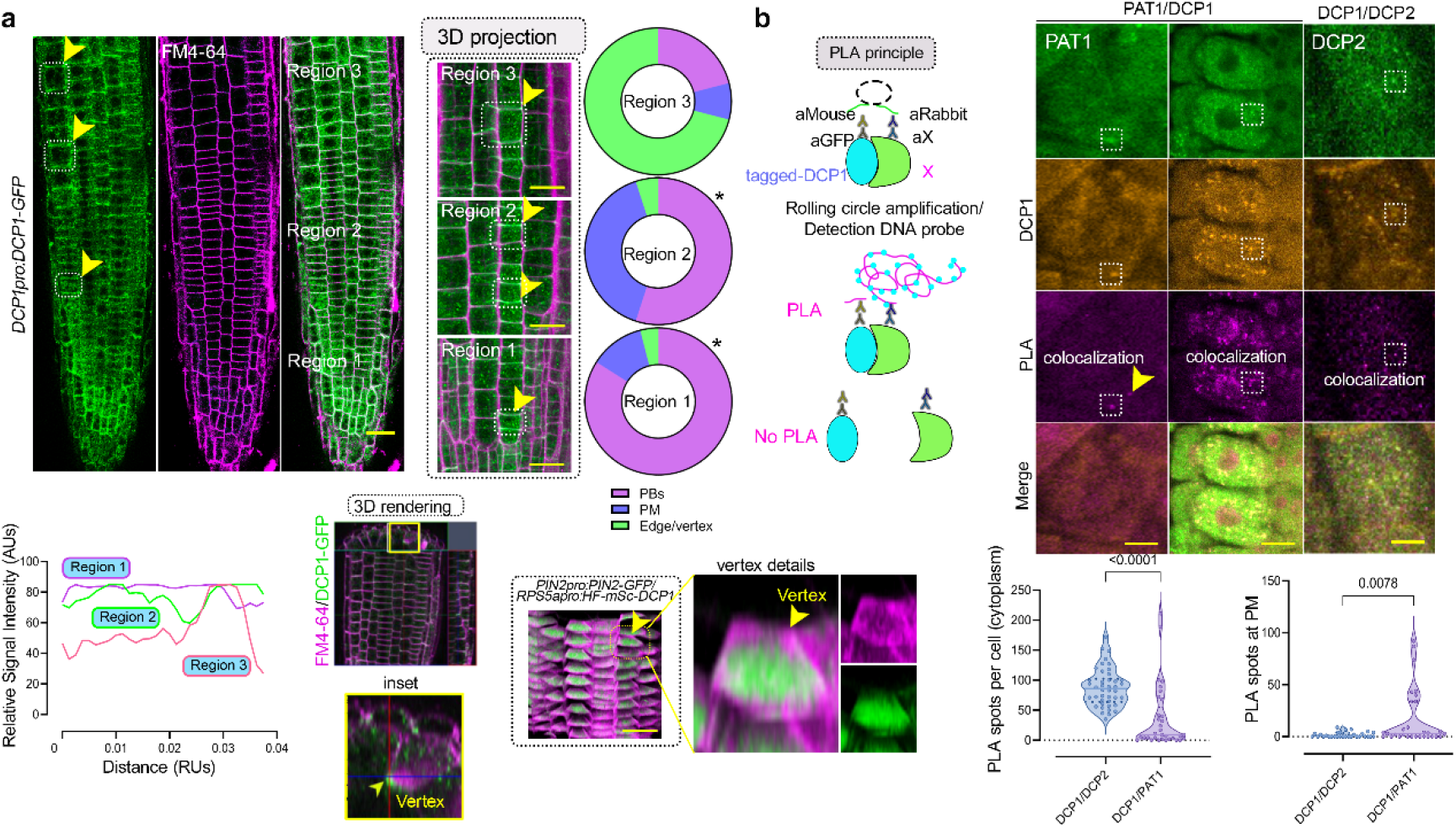
DCP1 protein accumulates at vertices during development. **a** DCP1 accumulates at edges/vertices. Right: three-dimensional (3D) projection of DCP1-GFP (green) in the whole root compared to FM4-64 staining (magenta, staining membrane/endosomes). The circular plots indicate DCP1 localization from 3D micrographs (regions 1–3; *N*=3, *n*=10 cells; asterisk: significant in a nested *t*-test at *p*<0.05, comparing regions 1–2 and 2–3). Bottom: DCP1 signal intensity among the three different regions, at the edge/vertex. Scale bars, 50 μm. Right: 3D-rendered images showing the localization of mScarlet-DCP1 at edges/vertices (in comparison to PIN2 signal). Scale bar, 20 μm. Arrowhead denotes a DCP1-positive vertex. **b** DCP1/DCP2 do not interact at edges. PLA assays for the confirmation of interactions (*in situ* proxitomes at distances < 40 nm). Left: principle of the method (see also text). Duolink® Proximity Ligation Assay (PLA) allows *in situ* detection of endogenous protein interactions with high specificity and sensitivity. Protein targets can be readily detected and localized with single-molecule resolution. Typically, two primary or antibodies against epitope tags raised in different species are used to detect two unique protein targets. A pair of oligonucleotide-labeled secondary antibodies (PLA probes) bind to the primary antibodies. Next, hybridizing connector oligos join the PLA probes only if they are near each other, and ligase forms a closed, circle DNA template that is required for rolling-circle amplification (RCA). The PLA probe then acts as a primer for a DNA polymerase, which generates concatemeric sequences during RCA. This allows up to a 1,000-fold signal amplification that is still tethered to the PLA probe, allowing localization of the signal. Last, labeled oligos hybridize to the complementary sequences within the amplicon, which are then visualized and quantified as discrete spots (PLA signals). Right: single optical sections from micrographs showing PLA-positive signal (resembling spots) produced by anti-FLAG/anti-GFP (*HF-mScarlet-DCP1*, *PAT1pro:PAT1-GFP* and *35S:DCP2-YFP*). Scale bars, 5 μm. Bottom: positive PLA spots per cell or on the PM (*N*=3, *n*=16–33 cells, significant difference determined by ordinary one-way ANOVA). The arrowhead in PAT1 PLA denotes a condensate. As a cautionary technical note, the “spots” do not connote physiologically relevant puncta. The colocalization of PLA spots with signals is also shown (colocalization).

We tested whether DCP1 accumulation at the PM or the edge/vertex was linked to localized decapping activity, which might delineate a spatially restricted decapping-active site, perhaps for localized regulation of translation. Ultra-fast live-cell imaging super-resolution microscopy showed that DCP1-decorated PBs undergo fission mostly at the PM plane, probably to release DCP1 from PBs (compare Fig. 3b versus Supplementary Fig. 1b). Likewise, in animals, PBs undergo fission at the ER membrane^16^. The decapping complex comprises DCP1 and DCP2 as minimal components^24^, but DCP1-GFP accumulation in edges was inversely correlated with its localization at PBs (Fig. 3c). Vertex localization was also unaffected by CHX which desolves decapping complexes (Supplementary Fig. 8b, arrowhead)^19^. These results argued against the existence of an active decapping complex at the edge/vertex. Furthermore, when we tested the dynamic interaction between DCP1 and DCP2, as a proxy for decapping activity (*sensu stricto* DCP1–DCP2 complex formation), by sensitized emission Förster resonance energy transfer (SE-FRET), we did not detect DCP1–DCP2 complex formation at the vertex (or the PM; Supplementary Fig. 9a). However, DCP1 did show a weak interaction with Protein Associated with Topoisomerase II 1 (PAT1) at the PM but not at the edge, where PAT1 does not accumulate (Supplementary Fig. 9a). This finding was consistent with the fact that at least some PBs first associate with the PM as an entity, before releasing their components. Together, these analyses speak against the existence of a localized decapping activity at the PM, although we cannot rule out the possibility that DCP1 transports RNA molecules to this site for localized translation.

To validate the lack of stable association between DCP1 and DCP2 or PAT1 at membranes, as FRET is sensitive to stoichiometry and may fail to capture very transient interactions, we exploited the power of quantitative 3D proximity ligation assays (PLAs^14, 25^). PLA uses complementary oligonucleotides fused to antibodies to determine the frequency with which proteins of interest find themselves nearby. Hence, PLA is highly relevant to the *in situ* “proxitome” and may not depend on direct interactions. DCP1 and DCP2 interacted only in the cytoplasm but not at the PM (Fig. 4b). We also tested DCP1 against PAT1 (positive control) and PIN7 (negative control) to assess the reliability of the assay in Arabidopsis roots, with the prediction that fewer interactions should occur when target protein pairs have increasingly discrete accumulation domains (i.e., PIN7; absent in APEAL, Supplementary Fig. 9b,c). Unexpectedly, through this approach, we established that DCP1 and DCP2 form complexes also in the nucleus (Supplementary Fig. 9b, insets), as was shown in yeast (*Saccharomyces cerevisiae*), where they act as a decapping reservoir^26^. This finding confirmed that PLA in this context can offer the required sensitivity to detect *in situ* proxitomes, providing additional informaton of interacting sites. Taken together, our results suggested a dynamic competition between PBs and vertices exclusively for the DCP1 component, irrespective of decapping.

### SCAR–WAVE recruits DCP1 at the cell edge/vertex

The accumulation of DCP1 during development at cell edges/vertices prompted us to examine the underlying mechanism by which DCP1 is recruited there. To this end, we examined DCP1 localization along the developmental root axis in more detail and compared this to that of other proteins that localize at cell edges/vertices and are enriched in the PDL step from subnetworks 3 and 4 (i.e., SOSEKI3 [SOK3], which regulates a cellular coordinate geometric system^27^ with log_2_FC =0.58 for NS and 2.0 for HS; and the SCAR–WAVE complex^28^ that regulates actin nucleation). In addition, as a negative control, we used the edge-localizing Ras-related protein Rab-A5c^29^, which we did not identify by APEAL. SOK3 localized at the cell division zone (the cell plate fusion site), unlike DCP1, which was absent from this site (Supplementary Fig. 10a,b). Later in development, SOK3 showed an almost perfect signal colinearity with DCP1 at edges/vertices. In contrast, Rab-A5c showed little colocalization with DCP1 at all stages examined (Supplementary Fig. 10c). We further observed that neither the loss of SOK3 function (from a *sok3* mutant), the simultaneous loss of SOK1/SOK3 functions (from genome editing of *SOK1* in the *sok3* mutant; details in Methods), nor Rab-A5c depletion (through a dominant-negative dexamethasone-inducible expression of inactive Rab-A5c^22^) resulted in changes in DCP1 localization at the PM or the edge/vertex (Supplementary Fig. 10b,c). These data suggested that neither SOK3 nor Rab-A5c are the initial recruiters of DCP1 at vertices/edges.

In sharp contrast, we observed an almost perfect signal colinearity at vertices/edges for mCherry and GFP signals in cells co-expressing *SCAR2-mCherry* and *DCP1-GFP* or *BRK1-YFP* (components of the SCAR–WAVE complex; ref^28^) under both NS and HS conditions (Fig. 5a). Of note, other SCAR–WAVE components like PIR121/SRA1 and GRL/NAP125 were highly enriched in PDL datasets, especially upon HS (log_2_FC ≥ 2 for NS and 3.5 for HS; see also Supplementary Fig. 7 and Supplementary File 7). Accordingly, upon HS, the PB numbers increase and the decrease of DCP1 association of DCP1 with the PM, were followed by a stronger colocalization at edges/vertices between SCAR2 and DCP1 (Fig. 3e and 5a). Hence, these results explained the APEAL results well predicting stronger association of DCP1–SCAR–WAVE.

**Fig. 5.**
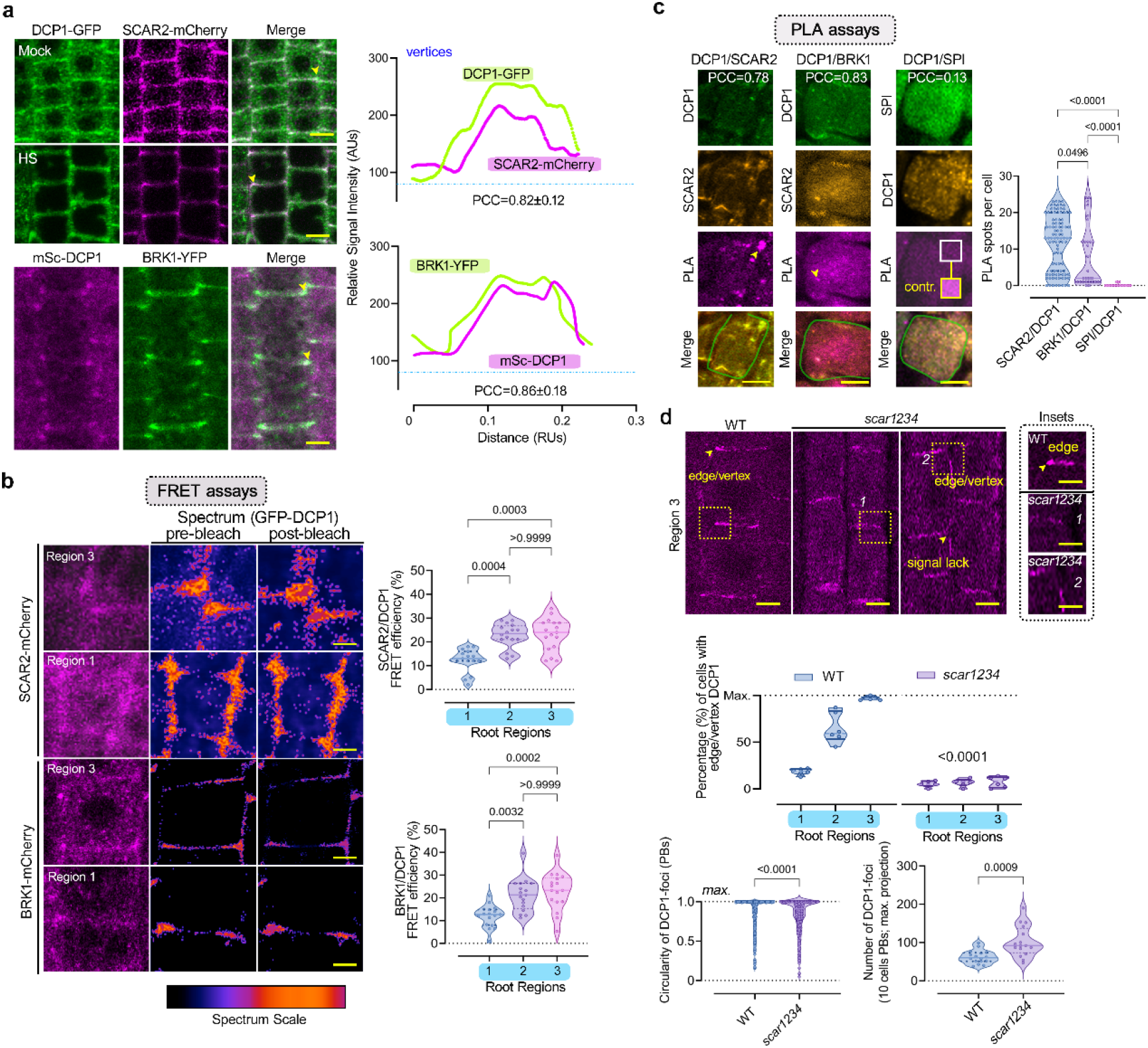
DCP1 cooperates with the SCAR–WAVE complex. **a** DCP1 colocalizes with the SCAR–WAVE components SCAR2 and BRK1. Representative confocal micrographs showing colocalization between DCP1 and SCAR2 or DCP1 and BRK1 (in lines coexpressing *DCP1pro:DCP1-GFP* and *SCAR2pro:SCAR2-mCherry* or *RPS5apro:HF-mScarlet-DCP1* and *BRK1pro:BRK1-YFP*) in root epidermal or cortex cells (regions 2–3). Right: relative signal intensity profiles of DCP1 or SCAR2 at vertices. Colocalization at these regions was also calculated as a Pearson’s correlation coefficient (PCC; *N*=2, *n*=4–10 edges/vertices in regions 2–3). Scale bars, 10 μm. **b** Representative confocal micrographs with acceptor photobleaching-FRET efficiency between SCAR2-mCherry (up) or BRK1-mCherry (down) and DCP1-GFP (epidermal cells, regions 1 and 3). Scale bars, 5 μm. Right: normalized FRET efficiency between SCAR2 or BRK1 and DCP1, respectively, among the different developmental root regions (*N*=2, *n*=16, significant difference determined by ordinary one-way ANOVA). **c** Representative confocal micrographs showing PLA spots produced by anti-GFP/anti-RFP in *DCP1-GFP/SCAR2-mCherry* lines, anti-FLAG/anti-GFP in *HF-mScarlet-DCP/BRK1-YFP* lines and *HF-mScarlet-DCP1/SPI-YPet* (SPIRRIG [SPI] is a negative control as it localizes to PBs only during salt stress and was not found in the APEAL). In the SPI PLA, a high-contrast inset is presented. Scale bars, 10 μm. Right: number of positive PLA spots per cell (*N*=3, *n*=16–33 cells, significant difference determined by one-way ANOVA). In the merged images, the cell contours are shown. **d** Representative confocal micrographs showing aDCP1 localization in the wild type (WT) or the *scar1 scar2 scar3 scar4* (*scar1234*) quadruple mutant (region 3). Scale bars, 10 μm. The arrowheads denote the lack of robust DCP1 localization in *scar1234* at the edge/vertex. Small panels at right show details corresponding to the regions delineated by dashed lines; scale bars, 1 μm. Bottom: quantification of PB numbers (DCP1-positive foci), percentage of cells with proper edge/vertex localization (*N*=3, *n*=16–33 cells, significant difference determined by ordinary one-way ANOVA) and DCP1-positive puncta circularity in region 3 for the WT and *scar1234* (*N*=1, *n*=1,610; significant difference determined by Kruskal-Wallis).

We also investigated whether the colocalization between DCP1 and SCAR–WAVE reflected an *in vivo* interaction. Indeed, DCP1 and SCAR2 or DCP1 and BRK1 interacted using FRET assays; the interaction took place mainly at edges/vertices but not in PBs (Fig. 5b). To ascertain whether DCP1 and SCAR–WAVE interacted, we estimated the strength of these interactions using quantitative 3D PLA (Fig. 5c). PLA determined that DCP1 and SCAR–WAVE components mainly interact at edges/vertices, although we occasionally observed PLA spots in the cytoplasm. We note here that APEAL failed to retain the interaction between DCP1 and SCAR–WAVE, indicating that co-immunoprecipitation experiments are ineffective (Supplementary File 7). DCP1 failed to produce positive PLA spots when combined with SOK3 or SPIRRIG (a protein that associates with SCAR–WAVE in root hairs or PBs only during salt stress; ref^30^(Fig. 5c and Supplementary Fig. 10a).

Furthermore, in the quadruple loss-of-function mutant *scar1234* that lacks the SCAR complex and its activity^31^, DCP1 localization at the edge/vertex was almost completely lost (Fig. 5d). In *scar1234* cells that had remnants of DCP1 at the edge, the DCP1 localization domain was expanded (Fig. 5d), but it still localized to the PM, suggesting that this localization does not require SCAR components. Furthermore, this reduced DCP1 localization at edges/vertices was associated with an increase in the number of DCP1-positive PBs and reduced circularity (Fig. 5d), suggesting that PBs in *scar1234* harden with time.

SCAR–WAVE also activates the Arp2–Arp3 complex (Actin-Related Protein 2 and 3 complex) to nucleate actin^32^; Arp2–Arp3 complex components were highly enriched in the APEAL datasets, again mainly under HS (Log_2_FC ∼1 for NS and 1.8 for HS; Supplementary Fig. 7, Supplementary File 7). The Arp2–Arp3 complex regulates the initiation of actin polymerization and the organization of filaments into y-branched networks. The loss of Arp2–Arp3 function (in the “crooked” mutant allele, *arpc5*^33^) did not deplete DCP1 from the vertex/edge, but rather led to a more variable DCP1 localization domain size at edges/vertices (Supplementary Fig. 10d). These results suggested that SCAR–WAVE, but not the effector Arp2–Arp3, may be involved in the recruitment of DCP1 molecules at the vertex.

### DCP1 phosphostatus defines its interaction with SCAR–WAVE and regulates actin

We tested whether DCP1 might affect actin organization at edges/vertices by modulating DCP1 localization. DCP1 showed an almost perfect signal colinearity with cortical F-actin (decorated by the *LifeAct-mCherry* marker expressed under the *UBIQUITIN 10* [*UBQ10*] promoter^34^) at edges/vertices (Fig. 6a; top). Furthermore, the actin-depolymerizing drug cytochalasin D enhanced DCP1 localization at edges/vertices (Fig. 6c), suggesting that actin does not recruit DCP1 there (or Arp2–Arp3, as discussed above and in Supplementary Fig. 10d). In contrast, treatment with the microtubule (MT)-depolymerizing drug amiprophos-methyl (APM)^35^ led to a more variable DCP1 domain size at the edge/vertex, in a site that would otherwise be likely confined by MTs (Supplementary Fig. 11a-d). Although MTs affected DCP1 confinement at edges/vertices, we opted against using MTs as a tool to affect DCP1 localization, as this would lead to pleiotropic effects. We thus looked for another finer approach to affect DCP1 localization at the cell edge. As the phosphorylation of residue Ser237 of DCP1 modulates its localization in PBs^36^, we asked whether DCP1 phosphorylation status might also modulate DCP1 abundance at the edges/vertices. We thus introduced a construct expressing the nonphosphorylatable variant *DCP1^S237A^-GFP* in *dcp1-1*^36^, a stronger allele than *dcp1-3*^18^, under the control of the *DCP1* promoter. We noticed an increased accumulation of fluorescence at edges/vertices in the resulting transgenic plants, compared to *DCP1pro*:*DCP1-GFP* in the *dcp1-1* (Fig. 6b). Conversely, introduction of the phosphomimetic variant *DCP1^S237D^-GFP* in *dcp1-1* showed a prevalent localization to PBs, alongside a pronounced inability to localize in a timely fashion to cell edges/vertices (Fig. 6b). In the lines expressing *DCP1^S237D^*, F-actin largely failed to accumulate at the cell edges/vertices (Fig. 6a; but not on PM), whereas *DCP1^S237A^* exerted the opposite effect, enhancing actin restriction at the edge/vertex (Fig. 6a). We confirmed that the *scar1234*, *dcp1-1* and *dcp1-3* mutants all display a similar lack of actin accumulation at edges/vertices, further demonstrating that actin restriction at edges/vertices increases along the developmental root axis (Fig. 6d, Supplementary Fig. 11d; comparison of region 1 vs 3). Altogether, these data establish that the SCAR–WAVE-DCP1 pathway controls actin at edges/vertices in a process modulated by the phosphorylation status of DCP1 Ser237.

**Fig. 6.**
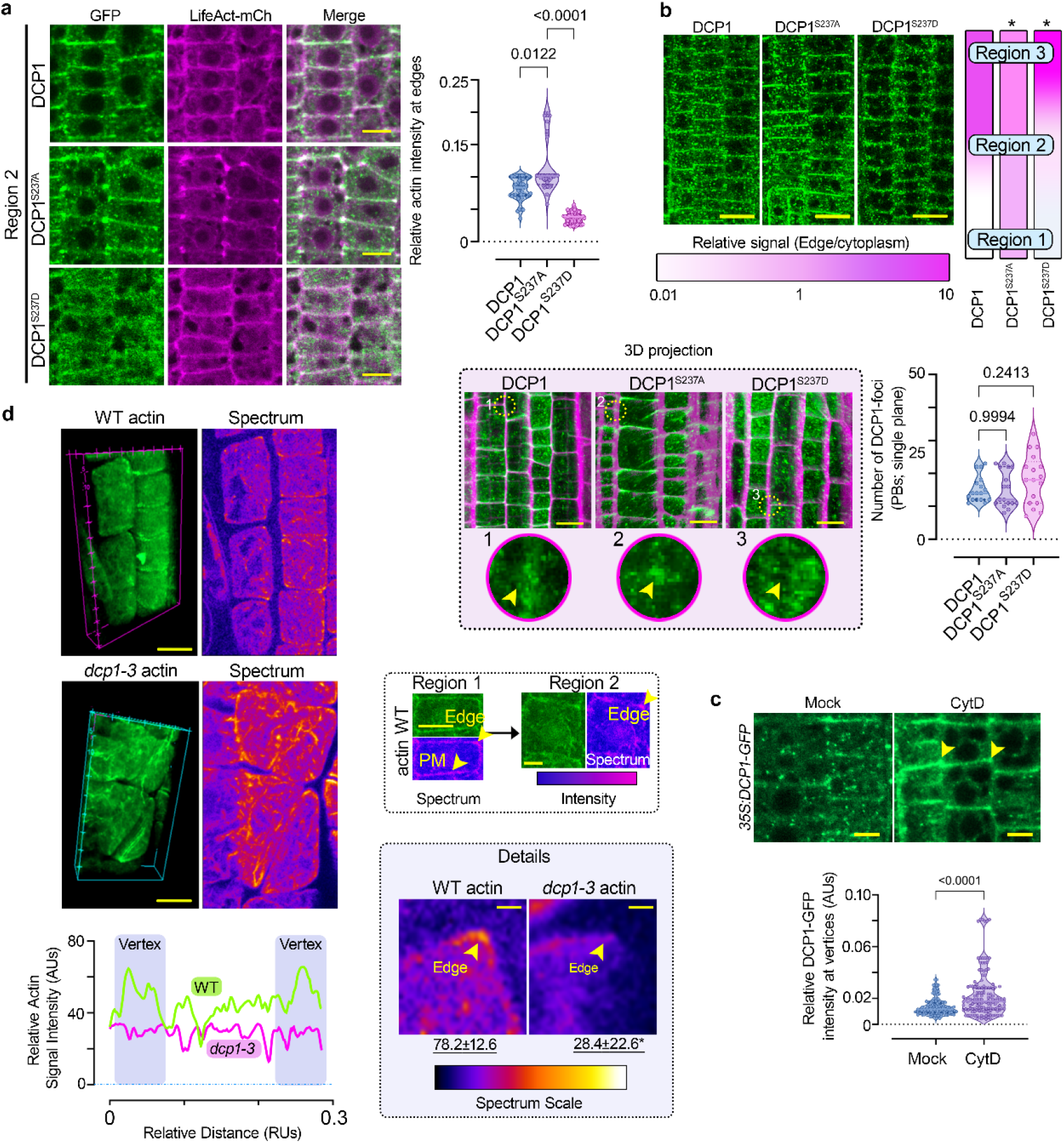
DCP1 phosphorylation regulates its interaction with SCAR–WAVE and actin accumulation at the edge/vertex. **a** Representative confocal micrographs showing colocalization between DCP1 or two DCP1 phosphovariants with LifeAct-mCherry in lines co-expressing *GFP-DCP1* (or variants) and *LifeAct-mCherry* (driven by the *DCP1* or *UBQ10* promoters, respectively). Scale bars, 7 μm. Right: relative signal intensity of actin at cell edges/vertices in epidermal cells (*N*=3, *n*=20–30, regions 2–3, significant difference determined by Kruskal-Wallis). **b** Representative confocal micrographs showing the effect of cytochalasin D (Cyt D; actin-depolymerizing drug) treatment on DCP1 localization (*N*=3). SCAR–WAVE increases at the PM in animals upon treatment with actin-depolymerizing drugs^56^. Note that Cyt D reduces DCP1-positive PBs. Arrowheads denote the edge/vertex. Bottom: relative signal intensity of DCP1-GFP in edges/vertices upon CytD treatment (*N*=2, *n*=129–140 edges or vertices, regions 2–3 averaged, significant difference determined by Wilcoxon). Scale bars, 4.5 μm. **c** Representative high-resolution confocal micrographs showing the localization of DCP1-GFP (or variants, region 2). Scale bars, 20 μm. Note the lack of isotropic growth in *DCP1^S237D^* (see also Fig. 6 and relevant discussion in the text). Middle: signal at the vertex, expressed as a color-coded edge/cytoplasmic signal ratio; asterisks indicate significant differences relative to intact DCP1 at *p*≤0.005, significant difference determined by ordinary one-way ANOVA. Right: relative signal intensity of actin at cell edges/vertices in epidermal cells (*N*=3, *n*=20–30, region 2–3, significant difference determined by ordinary one-way ANOVA). Bottom: 3D projection from super-resolution (120 nm axial) images of root meristematic cells captured from *DCP1pro:DCP1-GFP* and phosphovariants. Circular insets show the differential vertex localization of DCP1 (absent in *DCP1^S237D^*), and arrowheads denote the vertex. Scale bars, 10 μm. Note the enhanced accumulation of DCP1^S237A^-GFP at the vertex. **d** Representative confocal 3D rendering micrographs of root meristematic cells from the WT or the *dcp1-3* mutant immunostained with phalloidin for actin visualization. The “Spectrum” micrographs indicate the maximum color-coded signal intensity (scale on the right, middle inset). Scale bars, 2 μm (*z*-scale is 4 μm). Right: a detail of the higher actin accumulation at edges/vertices in region 2 (compare regions 1 and 2). Note that the signal is evenly distributed in region 1, whereas it mostly accumulates at the edge or vertex in regions 2 and 3 (see arrowheads; images on top). Insets (details) indicate the loss of vertex actin accumulation in *dcp1-3*. The arrowhead indicates the maximum color-coded signal intensity (“spectrum”: middle). Scale bars, 25 μm. Bottom: plot profile from the actin signal in the WT or *dcp1-3*. The two vertices are indicated.

### DCP1 solidification stabilizes SCAR-WAVE and actin at the vertex

The SCAR–WAVE/Arp2–ARP3 actin-filament-nucleating module specifies leaf pavement cell shape and trichome development, light-dependent and auxin-dependent root growth, stomatal opening, gravitropism, salt stress responses and immunity (ref^31^ and references therein) (see also Supplementary Fig. 11e for leaf pavement shape abnormalities in *dcp1-1*). We thus explored the possible consequences of the SCAR–WAVE–DCP1 interaction. Accordingly, we conducted FRET assays (acceptor photobleaching approach and sensitized emission, SE) along the proximodistal root axis between SCAR2 and DCP1 to determine when and where DCP1 and SCAR–WAVE most interact (Figs. 5b and 7a). These assays showed that SCAR–WAVE–DCP1 interaction increases along the proximodistal root axis during development (Fig. 7a; 3 regions, arrowhead), suggesting that the extent of interaction increases at “mature” edges/vertices. Furthermore, and as expected due to the increased colocalization between DCP1^S237A^ and SCAR2, DCP1^S237A^-GFP exhibited increased FRET with SCAR2-mCherry, and a faster response to the developmental increment, unlike DCP1^S237D^ (Fig. 7a,c).

**Fig. 7.**
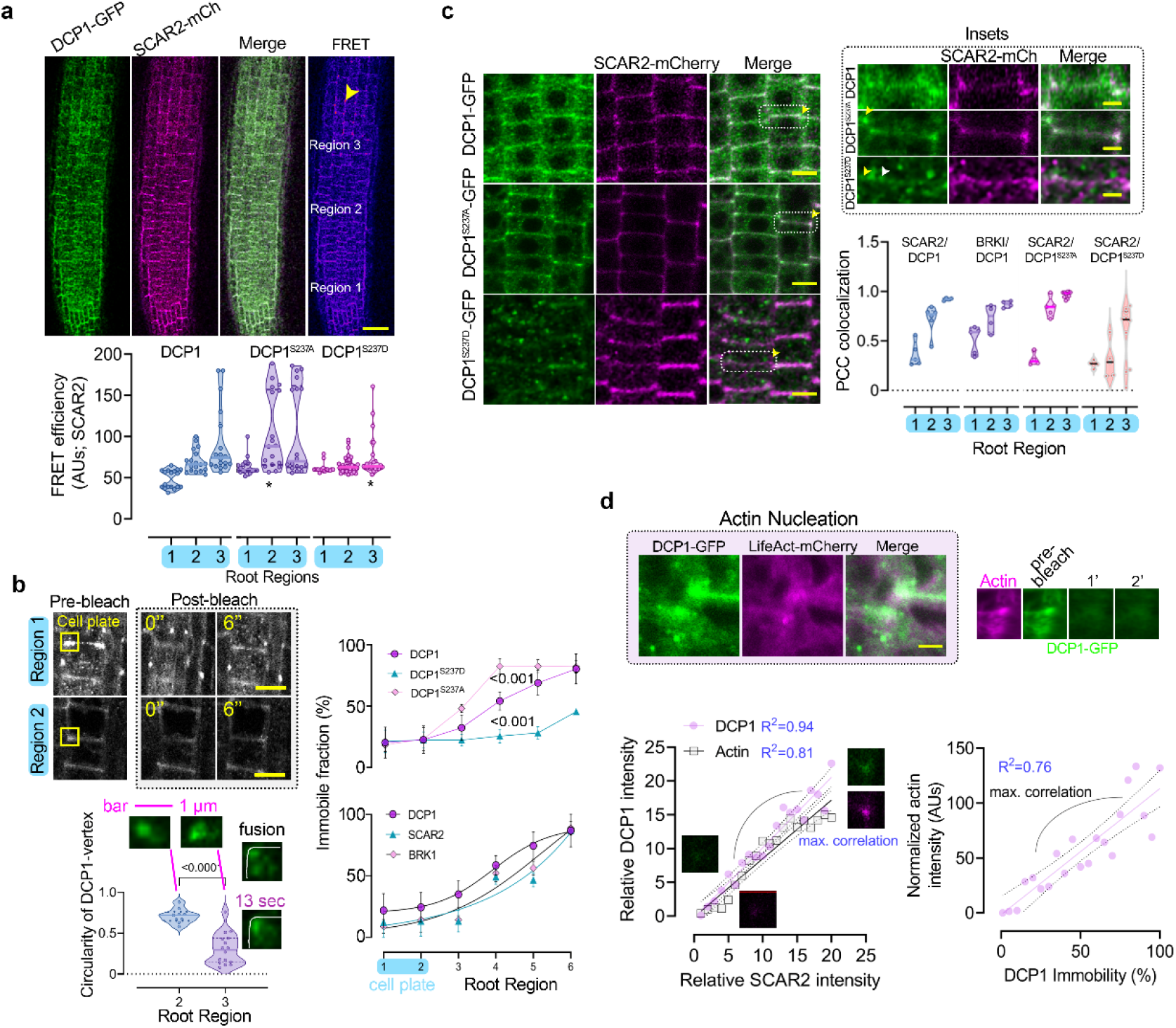
The liquidity of DCP1 defines interaction with SCAR-WAVE and actin nucleation. **a** SE-FRET efficiency between DCP1-GFP or its phosphovariants with SCAR2-mCherry (among the three different root regions; mainly epidermal cells). Scale bar, 50 μm. The arrowhead denotes high FRET efficiency at edges/vertices of region 3. Right: signal quantification of SE-FRET efficiency between the indicated combinations (*N*=6, *n*=10; \**p*<0.005 as determined by nested one-way ANOVA relative to the WT in the respective region). **b** Representative confocal micrographs showing FRAP assays from regions 1 and 2 for DCP1-GFP and analyses (below) showing DCP1-GFP (WT and phosphovariants), SCAR2-mCherry and BRK1-mCherry signal recovery and the immobile fraction at the PM. Data are means±SD (*N*=6, *n*=20; upper, nonlinear fit). Regions 1 and 2 correspond to immature and mature cell plates, respectively. In this instance, region 3 is split into TZ and DFZ (denoted as 5 and 6). Right: super-resolution microscopy calculations of DCP1 circularity in vertices (*N*=2, *n*=16, significant difference determined by Mann-Whitney). The insets show two examples of high and low circularity. A fusion event at the vertex is also shown (cell contours demarcated in white). **c** Representative confocal micrographs showing the colocalization between DCP1-GFP or phosphovariants and SCAR2-mCherry in root meristematic cells (region 2, epidermis). Scale bars, 10 μm. The insets at right show details of colocalization; the white arrow denotes the expansion of the SCAR2 domain. Scale bars, 1.5 μm. The graph indicates the relative signal intensity for the indicated combinations (as PCC; *N*=3, *n*=5). **d** Actin nucleation site, as indicated by DCP1-GFP and LifeAct localization. Right: correlation between immobile DCP1 fraction and DCP1/SCAR2 intensities or the correlation between DCP1/ACTIN (stained with anti-actin) and SCAR2 intensities (simple linear regression with the 95% confidence interval [CI] shown as dashed lines). The *R*^2^ values are also shown, along with representative micrographs (the FRAP shows a lack of signal recovery in this case).

We also noticed that DCP1 shows a gradual decline in total signal recovery (intensity) in FRAP assays at the vertex/edge during development (Fig. 7b and Supplementary Fig. 12a; ranging from 80 to 20% of the initial signal, *p*<0.05). Liquid-like protein assemblies can display rapid internal reorganization and a corresponding rapid fluorescent recovery on the order of seconds, whereas more solid-like condensates (formed by coarsening) may experience little recovery (mainly at their periphery)^14^. Hence, although not a decisive criterion itself (as discussed in the Supplementary Fig. 1 legend), this decrease in total signal recovery suggested that DCP1 can undergo solidification and coarsening, perhaps through a rise in local DCP1 protein concentration. Importantly, we observed a good correlation between DCP1 coarsening (judged here by the immobility in FRAP) and SCAR2–BRK1 or actin signal intensity/immobility at the vertex (Fig. 7b-d; correlation at the edge/vertex of LifeAct/DCP1 immobility *R*^2^∼0.75). Remarkably, while DCP1^S237D^-GFP could still localize at the edge/vertex (albeit later than the WT; ∼30 μm along the root axis, see also Fig. 5c quantifications), it failed to attain the immobile state and showed an expansion of the SCAR2 domain (Fig. 7b,c, insets). On the contrary, DCP1^S237A^-GFP showed a more rapid solidification than the DCP1-GFP. We confirmed these results using super-resolution imaging at cell edges, which revealed a decrease in the mean and robustness of circularity between regions 2 and 3 (means: 0.7 vs. 0.25; coefficient of variance: 12.59 vs. 62.55; *p*<0.0001). We also observed that DCP1 localizing at vertices in super-resolution images comprised multiple distinct puncta-like structures with the ability to undergo fusion, implying that DCP1 condensates form in a multilayered fashion (Fig. 7b, top right). Similarly, membrane-bound condensates in animals consist of multiple layers^4^, further supporting the idea that DCP1 exists in a phase-separated state at vertices. Hence, the material properties of DCP1 may affect SCAR–WAVE-actin nucleation.

### The SCAR–WAVE-DCP1 link can define growth anisotropy and developmental robustness irrespective of decapping

Edges are likely associated with uncharacterized trafficking pathways that may impinge on growth patterns and growth anisotropy^29^. Anisotropy, in terms of differential growth, is the relative change in principal dimensions over time, for example, the young hypocotyl elongates more than it widens^22^. SCAR–WAVE regulates growth patterns by impinging on cell wall properties at sharp cell edges (e.g., roots or trichome edge^28, 37^). We thus asked whether *dcp1-1* or *dcp1-3* or the phosphomimetic DCP1 variant showed similar phenotypic defects. To attenuate growth perturbations of SCAR mutants and focus on anisotropy, we used vertically grown plates with a high agar content, as described previously^38^. The roots of seedlings expressing *DCP1^S237D^*, of the progeny from *dcp1-1/DCP1* plants (as the homozygote cannot survive past the seedling stage, see also inset in Fig. 8a) and of the *dcp1-3* mutant, had slightly shorter roots than the WT (Fig. 8a).

**Fig. 8.**
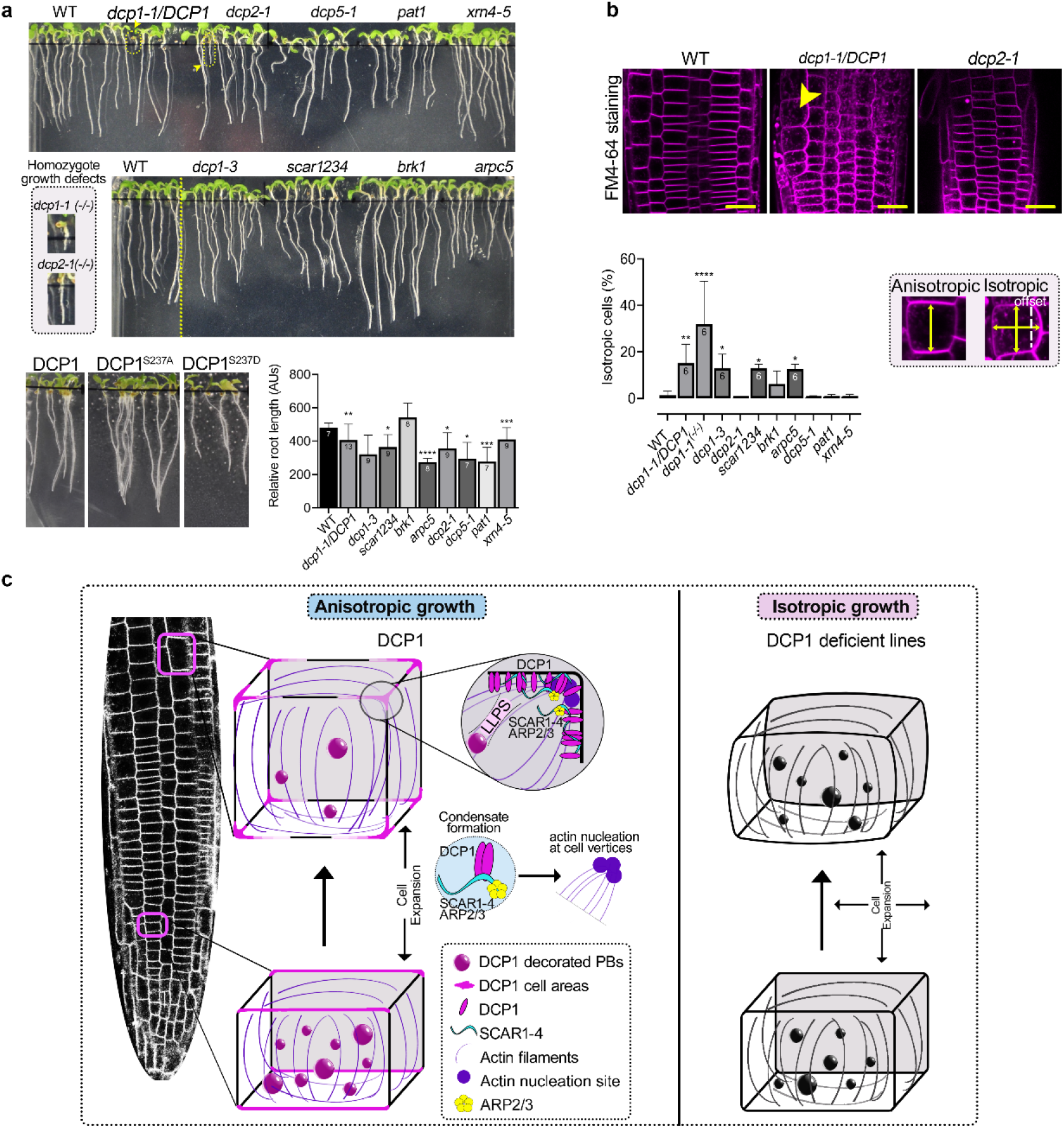
The SCAR–WAVE-actin-DCP1 nexus regulates development. **a** Images showing the phenotypes of *dcp1* mutants and mutants in PB components or SCAR–WAVE components. The insets show the growth defects of homozygous *dcp1-1* or *dcp2-1* mutants (denoted –/–). Bottom: relative root length of seedlings grown on 1.5% (w/v) agar plates. Data are means±SD (*N*=3, *n*=as indicated; significant difference determined by ordinary one-way ANOVA: \**p*<0.05; \*\**p*<0.01; \*\*\**p*<0.005; \*\*\*\**p*<0.0001). **b** DCP1 regulates cell expansion anisotropy. Representative confocal micrographs showing FM4-64 staining of the WT or *dcp1-1* or *dcp2-1* mutants. Scale bars, 20 μm. Bottom: percentage of isotropic cells per root meristem (%) in each genotype. Data are means±SD (*N*=3, *n*=as indicated; significant difference determined by ordinary one-way ANOVA). Examples of isotropic or anisotropic cells are shown, along with the developmental axis offset at the *x*- and *y*-axes. **c** Model for DCP1 function in root cells. Initially, DCP1 localizes at the cell plate (not the leading edge) and PBs and then occupies PM domains and PBs. Later, DCP1 accumulates more at the vertex, undergoing a phase transition during development that, upon cooperation with the SCAR–WAVE complex, helps actin nucleation at cell edges and mainly vertices.

Albeit with mild developmental defects, *dcp1-3* exhibited a significant loss of anisotropy similar to that seen in the *scar1234* or *arpc5* mutants, with fewer isotropic cells (Fig. 8a, b). We further asked if there was a correlation between the decapping machinery and the observed developmental defects. The greater anisotropy was independent of the presence of an active decapping complex, as *dcp2-1* (including the homozygous mutant, inset in Fig. 8a), *dcp5*, *xrn4* or *pat1* mutant seedlings lacked similar defects in anisotropy and actin, although they did exhibit reduced overall growth and plant stature (Supplementary Fig. 11d). However, seedlings expressing *DCP1^S237A^*had a longer root than WT seedlings, due to enhanced directional expansion (Fig. 8a). These results suggested that DCP1 and the SCAR–WAVE complex co-regulate anisotropy.

To consolidate the link between DCP1 and anisotropy, we used isoxaben, a cellulose biosynthesis inhibitor that promotes isotropic growth^29^. Long treatments with isoxaben increased the accumulation of DCP1-positive PBs (Supplementary Fig. 12b), so we determined a concentration and incubation time resulting in minimal effects on DCP1 localization for the following experiments. Isoxaben induced a marked reduction of anisotropic cell expansion in lines expressing *SCAR2-mCherry* and *DCP1pro:DCP1-GFP*, *DCP1^S237D^-GFP* or *DCP1^S237D^-GFP*, with a correlated signal diffusion of DCP1 and SCAR2 at cell edges (Supplementary Fig. 12c). Furthermore, isoxaben redistributed SCAR–WAVE and DCP1 in a similar and expanded domain before isotropy and radial growth (also known as “diffusible growth”; Supplementary Fig. 12b, inset). Notably, the DCP1^S237A^ cells expanded faster than those of the WT. If DCP1 did not have a function in isotropy, one would expect DCP1^S237A^ cells to simply swell more due to higher expansion propensity, which was not the case (Supplementary Fig. 12b). Because the enhanced feedback restriction of DCP1^S237A^–SCAR2 reduced radial growth, this result supports the notion that the SCAR–WAVE–DCP1 link regulates, drives and targets expansion anisotropy (Fig. 8c). Taken together, these results suggest that DCP1 and SCAR–WAVE are co-established at edges/vertices to participate in growth anisotropy.

## Discussion

The phase transitions of condensates necessitate a reevaluation of their cellular organization and how they impinge on physiology. We propose a framework describing how the composition of a cellular condensate (PBs here) can be determined and how we can integrate material information about condensates (liquid vs. solid state) to determine growth modules. We further exemplify here how a condensate is dissolved at the membrane and how a fraction of the condensate can be repurposed as a cellular coordinate geometric system. Unlike other coordinate systems like that driven by SOSEKI, the system suggested herein might be important later during development.

The plasticity of PBs allows a dynamic competition between condensates and membrane surfaces at vertices for the same components (e.g., DCP1). The subcellular positioning of DCP1 at certain membrane surfaces, and particularly at vertices, may be instrumental in driving phase transitions, by further reducing the radius within which proteins can diffuse, thereby promoting condensation and more rapid solidification through processes such as Ostwald ripening^39^. This mechanism further expands our comprehension of how condensation is promoted by reducing diffusion dimensionality (3D to 2D) from the cytoplasm to the plane of membranes. Likewise, dynamic spatial restriction influences the material states of condensates in neurons^40^.

Why is liquidity important at the onset of the PBs formation? Liquidity may allow greater equilibrium distinctions for binding to different partners, a necessity for building multiprotein complexes. Too much rigidity may also compromise the need for molecules to move during their function and could therefore impose kinetic barriers. Liquid-to-solid transitions of condensates depend on post-translational modifications, raising the intriguing possibility that a combination of vertex confinement for DCP1, together with DCP1 dephosphorylation, may entropically favor these transitions as shown here. Moreover, cell edges are sites of actin nucleation in plants (ref.^41^ and results herein), making the link between DCP1 and SCAR–WAVE highly relevant. In animal cells, LLPS increases the specific activity of molecules in the SCAR–WAVE pathway toward activating the Arp2–Arp3 complex^34^. There are also some thought-provoking parallels between plant edge/vertex condensation and animal epithelial sheets, where tight junctions are formed by ΖΟ condensates to maintain epithelial functions^1^; similarly, principles of actin nucleation in animals illustrate how LLPS can regulate stoichiometry in Arp2–Arp3 complexes^34^. Unlike other systems, in plants the situation is simpler, as Arp2–Arp3 appears to be activated solely by SCAR–WAVE, thus making plants an ideal system to study links between the SCAR–WAVE and Arp2–Arp3 complexes. Notably, the edge-decorating plant SOK proteins contain a PDZ domain that is also associated with tight junctions^1^. Notably, we did not find links to SOKs, whose exact functions remain to be determined.

In animal cells, many actin-nucleation-promoting factors, such as WHAMM (WASP homolog-associated protein with actin, membranes and microtubules), JMY (Junction Mediating And Regulatory Protein), the WASH complex and the SCAR–WAVE complex, can activate the Arp2–Arp3 complex^37^. Components of SCAR–WAVE undergo LLPS to promote actin nucleation. However, of the above list, only the SCAR–WAVE complex has been identified in plants thus far. Plants may thus employ other regulators to fulfill their needs for SCAR–WAVE condensation and actin nucleation. Vertices may therefore promote the activation of SCAR–WAVE through DCP1. We envision that this link may also remodel the transcriptional landscape, as the capture of DCP1 at vertices may reduce its potential to form PBs and thus store mRNAs poised for translation.

Perhaps the most puzzling contradiction in our data is the variable edge/vertex decoration by SCAR–WAVE or DCP1. The lack of robustness of this process may relate to stochastic condensation of SCAR–WAVE-DCP1 at regions closer to the QC, which can bring about local asymmetries in anisotropy at the cellular level. We accordingly show that cells with edges/vertices well defined by this complex follow a highly predictable anisotropic growth pattern, while cells with less determined SCAR–WAVE-DCP1 vertices have more diffusible growth patterns. Symmetry breakage, therefore, may entail randomized condensation that can bring about local asymmetries that underpin symmetries at the tissue level. Lastly, feedback between cell wall integrity and SCAR–WAVE-DCP1 may add more complexity to this system. We cannot discount links between SCAR–WAVE-DCP1 and the cell wall, in particular to vesicle trafficking and pectin, which can rigidify the middle lamella, the region between tricellular junctions (see also Fig. 2d, pectin, and Supplementary File 7).

## METHODS

### Plant materials

All the plant lines used in this study were in the Arabidopsis Columbia-0 (Col-0) accession except the ones indicated below. Primers used for genotyping and cloning are described in Table S1. Except for the T-DNA insertion mutants obtained from the NASC (The Nottingham Arabidopsis Stock Centre) as indicated in Table S1, the following mutants and transgenic lines used in this study were described previously: *dcp1-1*^17^, *dcp1-3*^18^, *dcp2-1*^42^, *dcp5-1*^43^, *pat1-1*^44^, *xrn4-5*^45^, *brk1*^28^, *scar1234*^28^, *arpc5*^46^, *35S:GFP-DCP1*^5^, *35S:DCP2-YFP*^47^, *dcp5-1 DCP5pro:DCP5-3HA*^43^, *RH12pro:RH12-GFP*^47^, *DCP1pro:DCP1-GFP*^48^, *DCP1pro:DCP1^S237A^-GFP*^48^, and *DCP1pro:DCP1^S237D^-GFP*^48^, *SOK3pro:SOK3-YFP*^27^, *SCAR2pro:mCherry-SCAR2*^49^, *BRK1pro:BRK1-mRuby*^28^, *BRK1pro:BRK1-YFP*^28^, *UBQ10pro;LifeAct-mCherry*^28^, *SPIpro:SPI-Ypet*^28^, *DEX˃RAB-A2c^DN^*^22^ and *RAB-A5cpro:RAB-A5c*^22^. In all experiments, plants from T1/F1 (co-localization experiments), T2/F2, or T3 (for physiological experiments) generations were used.

### Construction of the sok1sok3 CRISPR deletion mutant

EC1-driven *SOK1* deletion constructs were made as follows: single guide RNAs (sgRNAs) were synthesized with respective overhangs. The two complementary oligos were annealed and inserted into pEN-2xChimera using BpiI and BsmBI restriction enzymes. The sgRNAs were then transferred into pUbiCAS9-Red (for protoplasts) or pEciCAS9-Red (for stable transformation in *A. thaliana*) by Gateway® single-site LR recombination-mediated cloning. Efficiency of the sgRNAs was tested in protoplasts: Arabidopsis mesophyll protoplasts for transient expression of the CRISPR/Cas9 construct (pUbiCAS9-Red) were prepared as described previously with minor modifications^50^. Approximately 80,000 protoplasts were transformed with 16 μg of plasmid (pUbiCAS9-Red) and incubated for 48 h at 22°C under long photoperiod conditions (150 μmol m^−2^ s^−1^ and 16-h-light/8-h-dark cycles). Genomic DNA was isolated and concentrations adjusted before performing semi-quantitative PCR using oligonucleotides flanking the region targeted for deletion.

To create the *SOSEKI3* (At2g28150) deletion mutant, we used a previously described multiplexed editing approach^51^. We designed sgRNAs using the CRISPR-P 2.0 (http://crispr.hzau.edu.cn/CRISPR2)^52^ and CHOP-CHOP (https://chopchop.cbu.uib.no/)^53^ web tools. To delete the complete coding sequence of the gene, we targeted two sites near the start codon of *SOK3*, and one site downstream of the stop codon, along with equivalent sgRNAs for *SOK2* (At5g10150). Following the published protocol, we cloned all sgRNAs into a single pRU292 destination vector. This vector allows for zCas9i expression under the *UBi4.2* promoter, expression of the sgRNAs and has a FastGreen selection marker. sgRNAs and oligonucleotide sequences used for cloning are shown in Supplementary Table 1. The resulting binary vector was transformed into the *sok1* mutant background using the floral dipping method. We selected green-fluorescent T1 seeds and screened the resulting seedlings for large deletions in *SOK3* and absence of large deletions in *SOK2* using PCR (Supplementary Table 1). We validated the mutations in *SOK3* by direct gene-specific sequencing and crossed out the transgene by selecting non-fluorescent seeds in the T2 generation. We confirmed that *SOK2* was not deleted in this line, but there were small deletions in this gene. In the T3 generation, we selected double homozygous *sok1sok3* plants.

### Plant growth conditions

Arabidopsis seedlings were sterilized and germinated on half-strength Murashige and Skoog (MS) agar medium under long-day conditions (16 h light/8 h dark). In all experiments involving the use of mutants or pharmacological treatments, the medium was supplemented with 1% (w/v) sucrose or as otherwise specified. Arabidopsis plants for crosses, phenotyping of the above-ground part, and seed collection were grown on soil in a plant chamber at 22°C/19°C, 14-h-light/10-h-dark or 16-h-light/8-h-dark cycles, and light intensity 150 µE m^−2^ s^−1^. *Nicotiana benthamiana* plants were grown in Aralab or Percival cabinets at 22°C, 16-h-light/8-h-dark cycles, and a light intensity of 150 µE m^−2^ s^−1^.

### Phenotypic analysis and drug treatments

For quantification of phenotypes, seedlings were surface sterilized and grown on half-strength MS medium plates with 1% (w/v) sucrose. For a given genotype, differential contrast interference (DIC) images were captured on a Leica DM2500, Leica DM6B, or Leica DM6000. To define root length, images were captured from the plates using a Leica DM6000 with a motorized stage and computationally compiled together. Root length or size was determined using Image J/Fiji (National Institute of Health). For 1,6-hexanediol treatments, a 10% (v/v) aqueous solution was used. The stock solutions of 50 mM biotin, 10 mM cytochalasin D (Cyto D), 100 mM Amiprofos methyl (APM), 10 mM latrunculin B (Lat B), 50 mM brefeldin A (BFA) and 50 mM cycloheximide (CHX) were dissolved in DMSO, while 30 mM isoxaben was dissolved in ethanol. These inhibitors or drugs were diluted in half-strength MS medium with corresponding concentration and duration, and the final DMSO concentration was ≤0.1% (v/v) in all experimental analyses. Vertically grown 4- to 5-d-old Arabidopsis seedlings were incubated in half-strength liquid MS medium containing the corresponding drugs for each specific time course treatment as indicated.

### Bacterial strains, cloning and constructs

Electrocompetent Agrobacterium (*Agrobacterium tumefaciens*) strain C58C1 Rif^R^ (pMP90) or GV3101 Rif^R^ (i.e., a cured nopaline strain commonly used for infiltration) was used for electroporation and *Nicotiana benthamiana* infiltration. The Goldengate-compatible TurboID vector (35Spro:sGFP-TurboID-HF) was previously described ^6^. Cloning was done according to standard Goldengate cloning procedures. In brief, TurboID was synthesized with BsaI overhangs using the codon optimization tool of Integrated DNA Technologies for codon-optimized expression in Arabidopsis (Eurofins). The coding sequence of *DCP1* was PCR amplified using Phusion™ High-Fidelity DNA Polymerase & dNTP Mix (Thermo Fisher Scientific, Cat# F530N) using the GFP-DCP1 vector as a template. The 35S promoter carrying level 0 vector (pICSL13002) and all PCR parts with BsaI overhands were ligated to the Level1/2 vectors pICSL86900 and pICSL86922. Other constructs of *DCP1* were generated by Gateway cloning with pENTR-DCP1. For cloning, the bacterial strain NEB10 (New England Biolabs #C3019H) or NEB stable (New England Biolabs #C3040H; CRISPR constructs) was used. For BiFC and colocalization constructs, At1g33050, At2g26920, *ECT4*, *ECT6*, *MLN51*, *FLXL1*, *EIN2*, *VAP27-1* and At5g53330 were generated by Gateway cloning. pENTR vectors were generated via BP reaction with pDONR221 (Invitrogen) and PCR product amplicons from RT-PCR using cDNA from 1-week-old seedlings. As destination vectors, pSITE17 (nYFP-GW), pSITE18 (cYFP-GW) and custom-made pGWB601 with RFP were used.

### Immunoblotting

Infiltrated *N. benthamiana* leaves or Arabidopsis leaves and seedlings were harvested and their proteins extracted. The tissue samples were flash-frozen in liquid Ν_2_ and kept at –80°C until further processing. The samples were crushed using a liquid Ν_2_-cooled mortar and pestle, and the crushed material was transferred to a 1.5-mL or 15-mL tube. Extraction buffer (EB; 50 mM Tris-HCl pH 7.5, 150 mM NaCl, 10% [v/v] glycerol, 2 mM EDTA, 5 mM DTT, 1 mM phenylmethylsulfonyl fluoride, Protease Inhibitor Cocktail [Sigma-Aldrich, P9599] and 0.5 % [v/v] IGEPAL CA-630 [Sigma-Aldrich]) was added according to the plant material used. The lysates were pre-cleared by centrifugation at 16,000 × g at 4°C for 15 min, and the supernatant was transferred to a new 1.5-mL tube. This step was repeated two times and the protein concentration was determined by the RC DC Protein Assay Kit II (Bio-Rad, 5000122). Two times Laemmli buffer was added, and equivalent amounts of protein (∼30 μg) were separated by SDS-PAGE (1.0 mm thick 4 to 12% [w/v] gradient polyacrylamide Criterion Bio-Rad) in MOPS buffer (Bio-Rad) at 150 V. Subsequently, proteins were transferred onto a polyvinylidene fluoride (PVDF; Bio-Rad) membrane with 0.22-μm pore size. The membrane was blocked with 3% (w/v) BSA fraction V (Thermo Fisher Scientific) in phosphate buffered saline-Tween 20 (PBS-T) for 1 h at room temperature (RT), followed by incubation with horseradish peroxidase (HRP)-conjugated primary antibody for 2 h at RT (or primary antibody for 2 h at RT and corresponding secondary antibody for 2 h at RT). The following antibodies were used: streptavidin-HRP (Sigma-Aldrich; 1:25,000, N100), mouse anti-FLAG-HRP (Sigma-Aldrich, A8592, 1:2000), rat anti-tubulin (Santa Cruz Biotechnology, 1:1000), rabbit anti-GFP (Millipore, AB10145, 1:10,000), mouse anti-RFP (Agrisera, AS15 3028, 1:5,000), anti-mouse (Amersham ECL Mouse IgG, HRP-linked whole Ab [from sheep], NA931, 1:10,000), anti-rabbit (Amersham ECL Rabbit IgG, HRP-linked whole Ab [from donkey], NA934, 1:10,000), anti-rat (IRDye^®^ 800 CW Goat anti-Rat IgG [H + L], LI-COR, 925-32219, 1:10.000) and anti-rabbit (IRDye ^®^ 800 CW Goat anti-Rabbit IgG, LI-COR, 926-3221, 1:10,000). Chemiluminescence was detected with the ECL Prime Western Blotting Detection Reagent (Cytiva, GERPN2232) and SuperSignal™ West Femto Maximum Sensitivity Substrate (Thermo Fisher Scientific, 34094). The bands were visualized using an Odyssey infrared imaging system (LI-COR).

### APEAL approach details

After 24 h treatment (infiltration) with 50 μM biotin (diluted in 10 mM MgCl_2_, 10 mM MES pH 5.7), 2.5 mL pulverized Arabidopsis leaves or seedlings (∼0.5 g FW) were extracted in 5 mL EB (50 mM Tris-HCl pH 7.5, 150 mM NaCl, 10% [v/v] glycerol, 0.5 mM EDTA, 1 mM DTT, 0.5% [v/v] IGEPAL CA-630 [Sigma-Aldrich] and Protease inhibitor Cocktail [1:100 dilution, Sigma-Aldrich, P9599]). Extracts were incubated on a shaker at 4°C for 10 min and then centrifuged at 4°C, 13,000 g for 30 min. Five milliliters of clarified supernatants was incubated with 100 μL magnetic FLAG beads (Sigma-Aldrich, A8592) for 2 h at 4°C with gentle rotation, then the FLAG beads were precipitated with DynaMag™-2 Magnet (Thermo Fisher Scientific, 112321D) and washed four times with 1 mL EB (5 min each time). The supernatants (“flow-through1”) were filtered through PD-10 columns (Cytiva,17085101), then the biotin-depleted “flow-though2” was incubated with 100 μL Dynabeads™ M-280 Streptavidin (Thermo Fisher Scientific, 11205D) for 2 h at 4°C with gentle rotation. The FLAG beads and Dynabeads were washed five times with 1 mL EB and eight times with 1 mL 50 mM NH_4_HCO_3_ (5 min each time). The beads were then collected on a magnetic rack and subjected to on-beads digestion followed by mass spectrometry. For immunoblot analysis, the proteins were eluted from the FLAG beads with 60 μL 2x Laemmli buffer (Bio-Rad, 1610747) supplemented with 10 mM DTT and incubated at 95°C for 10 min. The proteins were eluted from the Dynabeads with 60 μL 2x Laemmli buffer supplemented with 5 mM biotin, 2% (w/v) SDS and 10 mM DTT and incubated at 95°C for 20 min.

### On-beads digestion

After immunoprecipitation and extensive washing with 25 mM NH_4_HCO_3_, 0.1 μg trypsin in 10 μL of 2 mM CaCl_2_, 10% (v/v) acetonitrile and 25 mM NH_4_HCO_3_ was added to beads and incubated at 37°C overnight. Fresh trypsin (0.1 μg) in 10 μL 25 mM NH_4_HCO_3_ was added to beads and incubated for another 4 h. The digested supernatant was transferred into a clean centrifuge tube, and acetonitrile was evaporated under a vacuum. The samples were then acidified and desalted using a C18 stage tip before being analyzed by nano-liquid chromatography-tandem mass spectrometry.

### Liquid chromatography-tandem mass spectrometry

Samples were analyzed by LC-MS using a Nano LC-MS/MS apparatus (Dionex Ultimate 3000 RLSCnano System) interfaced with an Eclipse Tribrid mass spectrometer (Thermo Fisher Scientific). Samples were loaded onto a fused silica trap column (Acclaim PepMap 100, 75 μm x 2 cm; Thermo Fisher Scientific). After washing for 5 min at 5 µL/min with 0.1% (v/v) trifluoroacetic acid (TFA), the trap column was brought in-line with an analytical column (Nanoease MZ peptide BEH C18, 130 A, 1.7 μm, 75 μm x 250 mm, Waters) for LC-MS/MS. Peptides were fractionated at 300 nL/min using a segmented linear gradient 4–15% B in 30 min (where A: 0.2% formic acid, and B: 0.16% formic acid, 80% acetonitrile), 15–25% B in 40 min, 25–50% B in 44 min, and 50–90% B in 11 min. Solution B was then returned to 4% for 5 min for the next run. The scan sequence began with an MS1 spectrum (Orbitrap analysis, resolution 120,000, scan range from m/z 350–1600, automatic gain control (AGC) target 1E6, maximum injection time 100 ms). The top S (3 sec) and dynamic exclusion of 60 sec were used for the selection of parent ions for MS/MS. Parent masses were isolated in the quadrupole with an isolation window of 1.4 m/z, automatic gain control (AGC) target 1E5, and fragmented with higher-energy collisional dissociation with a normalized collision energy of 30%. The fragments were scanned in Orbitrap with a resolution of 30,000. The MS/MS scan range was determined by the charge state of the parent ion but the lower limit was set at 100 amu.

### Database search

LC-MS/MS data were analyzed with Maxquant (version 1.6.10.43) with the Andromeda search engine. The type of LC-MS run was set to 1 (label-free). LC-MS data were searched against The Arabidopsis Information Resource (TAIR) plus a common contaminant database. Protease was set as trypsin, which allowed two mis-cuts. N-terminal acetylation and oxidation at methionine were set as variable modifications. Maximum two variable modification was allowed. iBAQ/LFQ values were also determined and used for protein validation. Protein and peptide false discovery rate (FDR) were set to 1%. Reverse hit and common contaminants, as well as proteins identified only by modified sites, were removed. GO term analysis was performed using a combination of the Panther database and BioConductor package in R.

### Visualization of networks and analyses

Cytoscape 3.5.1 was used. Tab-delimited files containing the input data were uploaded. Unless otherwise indicated, the default layout was an edge-weighted spring embedded layout, with NormSpec used as edge weight. Nodes were manually re-arranged from this layout to increase visibility and highlight specific proximity interactions. The layout was exported as a PDF and eventually converted to a .TIFF file.

### Agrobacterium-mediated transient transformation of Nicotiana benthamiana

*Nicotiana benthamiana* plants were grown under normal light and dark regimes at 25°C and 70% relative humidity. Three- to four-week-old *N. benthamiana* plants were watered from the bottom ∼2 h before infiltration. Transformed Agrobacterium strain C58C1 Rif^R^ (pMP90) or GV3101 Rif^R^ harboring the constructs of interest was used to infiltrate *N. benthamiana* leaves and for transient expression of binary constructs by Agrobacterium-mediated transient infiltration of lower epidermal leaf cells. Transformed Agrobacterium colonies were grown for ∼20 h in a shaking incubator (200 rpm) at 28°C in 5 mL of yeast extract broth (YEB) medium (5 g/L beef extract, 1 g/L yeast extract, 5 g/L peptone, 0.5 g/L MgCl_2_ and 15 g/L bacterial agar), supplemented with appropriate antibiotics (i.e., 100 g/L spectinomycin). After incubation, the bacterial culture was transferred to 15-mL Falcon tubes and centrifuged (10 min, 5000 g). The pellets were washed with 5 mL of infiltration buffer (10 mM MgCl_2_, 10 mM MES pH 5.7), and the final pellet was resuspended in infiltration buffer supplemented with 100 μM acetosyringone. The bacterial suspension was diluted with infiltration buffer to adjust the inoculum cell density to a final OD_600_ value of 0.2–1.0. The inoculum was incubated for 2 h at room temperature before infiltration into *N. benthamiana* leaves by gentle pressure infiltration of the lower epidermis of leaves (fourth and older true leaves were used, and about 4/5-1/1 of their full size) with a 1-mL hypodermic syringe without a needle.

### Bimolecular fluorescence complementation (BiFC)

The BiFC assay was done in *N. benthamiana* plants. Excitation wavelengths and emission filters were 514 nm/band-pass 530–550 nm for YFP, 561 nm/band-pass 600–630 nm for RFP and 488 nm/band-pass 650–710 nm for chloroplast auto-fluorescence. The objective used was a plan-apochromat 40x with NA=1.2 M27 (Zeiss).

### TIRFM imaging and tracking analyses

TIRF microscopy images were acquired using MetaMorph software on an Olympus IX-81 microscope. A DV2 image splitter (MAG Biosystems) was used to separate GFP and RFP emission signals. Time-lapse movies were obtained at 100-ms intervals. For MSD analysis, 30-s-long movies with 100-ms intervals and 200-ms exposure were used. Particle tracking was limited to the amount of time that plasma membrane remained at the focal plane; the median track length was 2,000 frames, corresponding to 3.5 s of imaging. The tracking of particles was performed with the Mosaic suite of Fiji or NanoTrackJ/TrackMate, using the following parameters: radius 3 of fluorescence intensity, a link range of 1, cutoff of 0.1%, and a maximum displacement of 8 px, assuming Brownian dynamics.

### Quantification of fluorescent intensity, FRAP and FRET

To create the most comparable lines to measure the fluorescence intensity of reporters in multiple mutant backgrounds, we crossed homozygous mutant plants carrying the marker with either a wild-type plant (to yield progeny heterozygous for the recessive mutant alleles and the reporter) or crossed to a mutant only plant (to yield progeny homozygous for the recessive mutant alleles and heterozygous for the reporter). Fluorescence was measured as a mean integrated density in regions of interest (ROIs) with the subtraction of the background (a proximal region that was unbleached and had less signal intensity than the signal of the ROI region). FRAP mode of Zeiss 780 ZEN software was set up for the acquisition of 3 pre-bleach images, 1 bleach scan, and 96 post-bleach scans (or more). Bleaching was performed using the 488-, 514- and 561-nm laser lines at 100% transmittance and 20–40 iterations depending on the region and the axial resolution (iterations increased in deeper tissues to compensate for the increased light scattering). In FRAP the width of the bleached ROI was set to 2–10 µm. Pre- and post-bleach scans were at minimum possible laser power (0.8% transmittance) for 458 nm or 514 nm (4.7%) and 5% for 561 nm; 512 x 512 8-bit pixel format; pinhole of 181 μm (>2 Airy units), and zoom factor of 2.0. The background values were subtracted from the fluorescence recovery values, and the resulting values were normalized by the first post-bleach time point and divided by the maximum fluorescent time-point set maximum intensity as 1. The objective used was a plan-apochromat 20x with NA=0.8 M27 (Zeiss). The following settings were used for photobleaching DCP1: 10–20 iterations for DCP1-GFP; 10 to 60 s per frame; 100% transmittance with the 458- to 514-nm laser lines of an argon laser. Pre- and post-bleach scans were at minimum possible laser power (1.4 to 20% transmittance) for the 488-nm and 0% for all other laser lines, 512 x 512 pixel format and zoom factor of 5.1. The fluorescence intensity recovery values were determined, the background values were subtracted from the fluorescence recovery values, and the resulting values were normalized against the first post-bleach time point. FRET analyses were conducted using the method described previously^14^.

### Immunocytochemistry, PLA and imaging

Immunocytochemistry was done as described previously ^54^. The primary antibody used was rabbit anti-PAT1 (diluted 1:500) ^44^, rat anti-tubulin YL1/2 (1:200; Santa Cruz Biotechnology), mouse anti-DCP1 (diluted 1:100; prepared herein) and anti-HA (diluted 1:300; Sigma-Aldrich). In brief, specimens were washed three times for 90 min in PBS-T and incubated overnight with anti-rabbit fluorescein isothiocyanate-conjugated (FITC) secondary antibody (Jackson ImmunoResearch, 711-095-152) diluted 1:200-250, DyLight™ 549 AffiniPure Fab Fragment Rabbit Anti-Mouse IgG (H+L) (Jackson ImmunoResearch, 315-507-003), Rhodamine Red™-X (RRX) AffiniPure Donkey Anti-Rabbit IgG (H+L) (Jackson ImmunoResearch, 711-295-152), Tetramethylrhodamine (TRITC) Donkey Anti-Rabbit IgG (H+L) (Jackson ImmunoResearch, 711-026-152) diluted 1:200–250. After washing in PBS-T and incubation with DAPI (1 μg/mL), specimens were mounted in Vectashield (Vector Laboratories) medium and observed within 48 h.

Immunostaining of actin was done as described previously with minor modifications^28^. Roots of 4-day-old Arabidopsis seedlings were fixed for 1 h in actin-fixation buffer (50 mM PIPES, pH 7.2, with 20 mM EGTA and 20 mM MgSO_4_ containing 2% [w/v] paraformaldehyde, 0.1% [v/v] Triton X-100 and 400 mM maleimidobenzoyl-N-hydroxysuccinimide ester [Thermo Fisher Scientific, 22311]); the cell wall was digested for 30 min (in 0.2% [w/v] driselase and 0.15% [w/v] macerozyme). Then samples were incubated for 30 min in blocking buffer containing 2% (w/v) BSA and then incubated with Alexa Fluor 488 phalloidin (1:400 dilution, Thermo Fisher Scientific, A12379) overnight at 4°C. After washing with PBST buffer three times, samples were mounted in Vectashield (Vector Laboratories, Burlingame, CA) and root tips were imaged using a Zeiss 780 confocal laser scanning microscope.

PLA immunolocalization was done as described previously ^14^. Primary antibody combinations diluted 1:200 for α-GFP mouse (Sigma-Aldrich, SAB2702197),1:200 for α-FLAG mouse (Sigma-Aldrich, F1804), 1:200 for α-RFP mouse (Agrisera, AS15 3028) and 1:200 for α-GFP rabbit (Millipore, AB10145) were used for overnight incubation at 4°C. Roots were then washed with MT-stabilizing buffer (MTSB: 50 mM PIPES, 5 mM EGTA, 2 mM MgSO_4_, 0.1% [v/v] Triton X-100) and incubated at 37°C for 3 h either with α-mouse plus and α-rabbit minus for PLA assay (Sigma-Aldrich, 681 Duolink). PLA samples were then washed with MTSB and incubated for 3 h at 37°C with ligase solution as described (Pasternak et al, 2018). Roots were then washed 2x with buffer A (Sigma-Aldrich, Duolink) and treated for 4 h at 37°C in a polymerase solution containing fluorescent nucleotides as described (Sigma-Aldrich, Duolink). Samples were then washed 2x with buffer B (Sigma-Aldrich, Duolink), with 1% (v/v) buffer B for another 5 min, and then the specimens were mounted in Vectashield (Vector Laboratories) medium.

### DCP1 antibody production

The *DCP1* cDNA in pGAT4 (hexahistidine-tagged vector) was transformed in BL21 (DE3) Rosetta or BL21 (DE3) Rosetta II *Escherichia coli* cells. Bacterial cultures were grown in 800 mL of Luria Bertani (LB) medium supplemented with 100 mg L^−1^ of ampicillin and 25 mg L^−1^ of chloramphenicol. Protein production was induced at OD_600_ = 0.5 with 0.05 to 1 mM isopropyl β-D-1-thiogalactopyranoside (IPTG). After 3 h, the cells were harvested by centrifugation at 2,500 g for 20 min at room temperature and frozen overnight at –80°C. Preparation of hexahistidine-tagged recombinant DCP1 was performed according to the manufacturer’s instructions in Tris buffer (Qiagen, 30210). The abundance of proteins was estimated by CBB staining by SDS-PAGE or on immunoblots using anti-his antibodies. The DCP1 protein was dialyzed overnight in assay buffer (2 L) and was used to immunize four mice. The antibodies were further purified by solid-phase absorption using columns with DCP1. Pre-immune sera were also collected and used as an additional negative control, producing no signal.

### Statistics

All statistical data show the mean ± SD of at least three biologically independent experiments or samples, or as otherwise stated. *N* denotes biological replicates, and “*n*” technical replicates or population size. Statistical analyses were performed in GraphPad (https://graphpad.com/) or R studio (R-project.org). Each data set was tested whether it followed normal distribution when *N*≥3 by using the Shapiro normality test. The significance threshold was set at *p*<0.05 (significance claim), and the exact values are shown in graphs. Graphs were generated by GraphPad Prism or R. Details of the statistical tests applied, including the choice of the statistical method, are indicated in the corresponding figure legend. In boxplots or violin plots, upper and lower box boundaries represent the first and third quantiles, respectively, horizontal lines mark the median and whiskers mark the highest and lowest values. Image analyses and intensity measurements (Integrated Density) were done using Fiji v. 1.52 software (rsb.info.nih.gov/ij). For relative values, the intensities were normalized against the background signal intensity in raw images/micrographs. The calculations were done with arbitrary units (denoted as AUs). The dwell time rate of tagged proteins in FRAP experiments was calculated by the single exponential fit^54^. Colocalization was analyzed using Pearson statistics (Spearman or Manders analyses produced similar results or trends)^55^. Images were prepared in Adobe Photoshop v. 2022. Time series movies were compressed, corrected and exported as .avi extension files. The nonspecific fluorescence decay was corrected using Fiji and default options using the bleaching correction tool. Videos were digitally enhanced with Fiji implemented filters, correcting noise using the Gaussian blur option and pixel width set to 1.0.

## Acknowledgments

We thank Sebastian Y. Bednarek, Morten Petersen, Ping He, Annie Marion-Poll, Shu-Hsing Wu, Jan Petrášek and Charlotte Kirchhelle for kindly providing published materials. This work was supported by the EU Marie Curie-RISE PANTHEON (P.N.M.), the Carl Tryggers Foundation (P.N.M.), and partly by the VR research council (P.N.M.), FORMAS research council (P.N.M.), Helge Axelsson Foundation (C.L.), IMBB-FORTH start-up funding (P.N.M.), European Research Council (ERC; ‘DIRNDL’, contract number 833867; D.W.) and Spanish grant PID2020-119737GA-I00 funded by MCIN/AEI/437 10.13039/501100011033 (E.G.B.).

## Author contributions

Conceptualization: P.N.M.; Methodology: C.L., A.M., A.M., A.V.; Investigation: C.L., A.M., A.V., E.G.B.; Visualization: C.L., A.M., P.N.M.; Funding acquisition: C.L., E.G.B., D.W., P.N.M.; Project administration: P.N.M.; Supervision: P.N.M.; Writing - original draft: P.N.M.; Writing - review & editing: C.L., A.M., E.G.B., A.V.

## Data availability

All data generated or analyzed during this study are included in this published article (and its supplementary information files). Source data are provided with this paper. Additional information and materials generated for and/or reported in this article are available from the corresponding author upon request. Source data are provided with this paper.

## Supplemental figures

**Supplementary Fig. 1.**
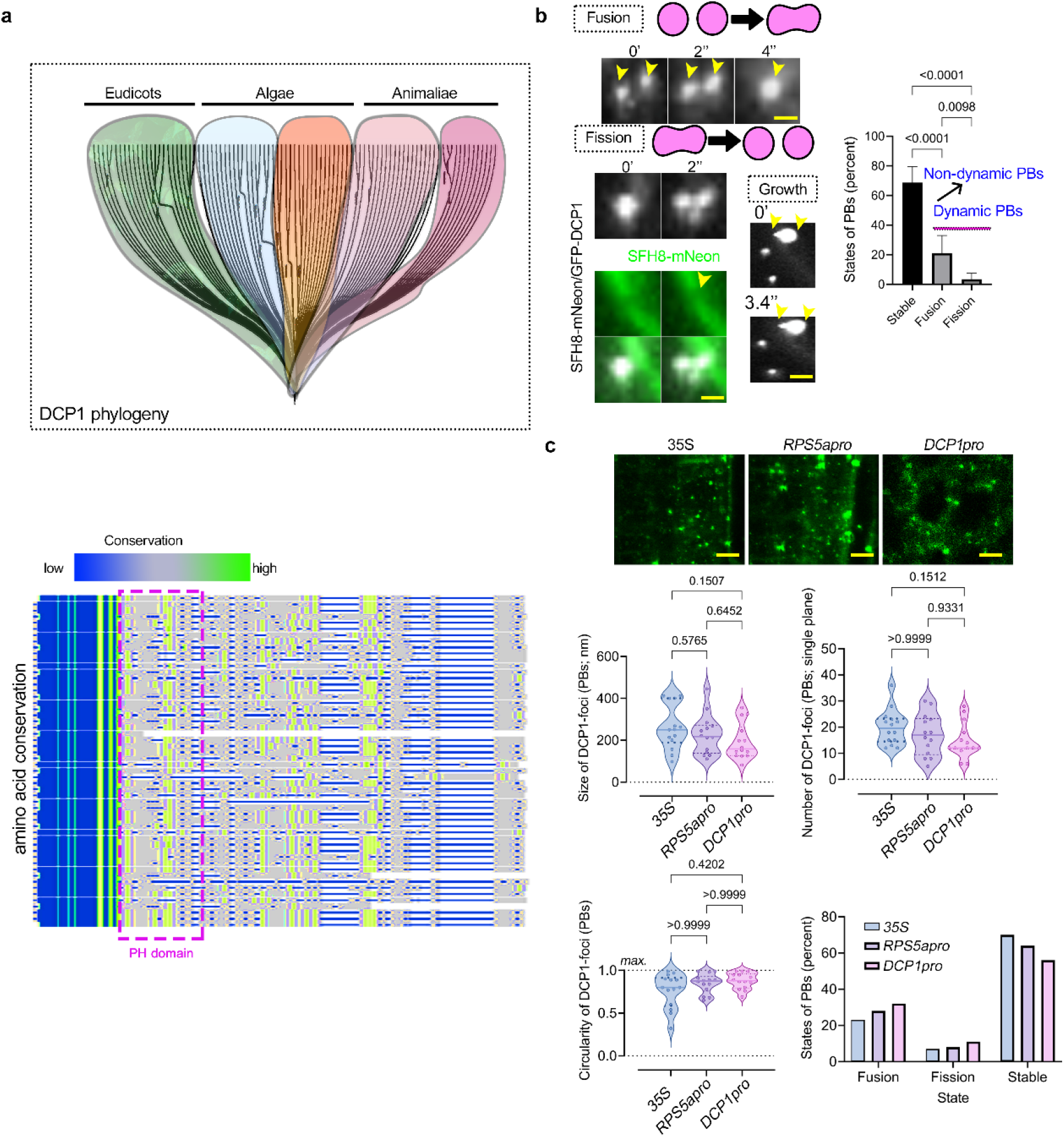
Phylogeny of DCP1 and dynamics of PBs in plants: DCP1- decorated PBs show LLPS features. **a** The DCP1 phylogeny (upper “umbrella phylogram”) is monophyletic. Bottom: conservation of amino acid residues. Note the conserved PH region (pleckstrin homology) in DCP1, which may allow DCP1 to interact with membrane lipids. Sequences used to build the phyllo- dendrogram are presented in Supplementary File 1. **b** Representative confocal images showing fusion (coarsening), fission and growth of PBs (DCP1-positive) decorated by DCP1 (in the *DCP1pro:DCP1-GFP* transgenic line) in root meristematic epidermal cells (proximal meristem zone 1, as defined later). Diagrams depict fusion and fission events. In “growth”, the two arrowheads denote the axis of growth in this particular punctum (note the offset of the lower arrow). Bottom: *DCP1-GFP* is co-expressed with *SFH8-mNeon* (decorating the PM; *SFH8pro:SFH8-mNeon*^14^). The arrowhead denotes a loss of SFH8 signal at the site where the fission takes place. Scale bars, 500 nm. Right: PB states in the cytoplasm (dynamic, fusion and fission; non-dynamic, stable; *N*=1, *n*=6–8; significant difference determined by ordinary one-way ANOVA). As a cautionary note, “stable” PBs may not show a dynamic behavior in the imaging time used (∼3–5 min) but may do so later. **c** Representative confocal images from epidermal root cells showing cytoplasmic and motile PBs decorated by DCP1-GFP derived from a transgene driving the expression of DCP1-GFP from the *35S*, *RPS5a* (strong meristematic promoter) or DCP1 promoter (two lines were checked per construct and produced similar results). Size, circularity (see also Supplementary Fig. 2 for an explanation of calculations), number (calculated manually from 2D micrographs of a single cell; we considered puncta above the diffraction limit of confocal microscopy, ∼200 nm) and states (fusion, fission and stable at the time-scale used, 3–5 min; *N*=1, *n*=6–8 [*N*=1, *n*=14 for size]) are indicated; significant difference determined by ordinary one-way ANOVA). Note the increased coarsening in the *35S*-driven lines, confirmed also by reduced fusion/fission implying increased viscosity and reduced elasticity. The *35S* and *RPS5a* promoters showed larger variance in PB sizes (CI lower % 201.1, 171.8 and 144.0). Yet, all cell types examined, irrespective of the promoter used, have DCP1-positive PBs that show circularity, fusion, fission and rapid fluorescence recovery after photobleaching (FRAP, *t*_1/2_=12.1±4.3 sec; meristematic epidermal cells). FRAP results imply that DCP1 molecules diffuse in and out of PBs and likely suggest that PBs are in an LLPS state. Circularity and the tendency to undergo fusion and fission, reflecting surface tension characteristics of a liquid, are good indicators of liquidity. FRAP, albeit a popular method to quantitatively evaluate the fluidity of droplets in cells, can occur via molecular diffusion, dissociation-binding or any kind of turnover in the target area (defined as the region of interest); thus, turnover observed by FRAP is not necessarily a consequence of fluidity. Hence, FRAP datasets should be accompanied by additional datasets, e.g., circularity and hexanediol treatment (see also Supplementary Fig. 2).

**Supplementary Fig. 2.**
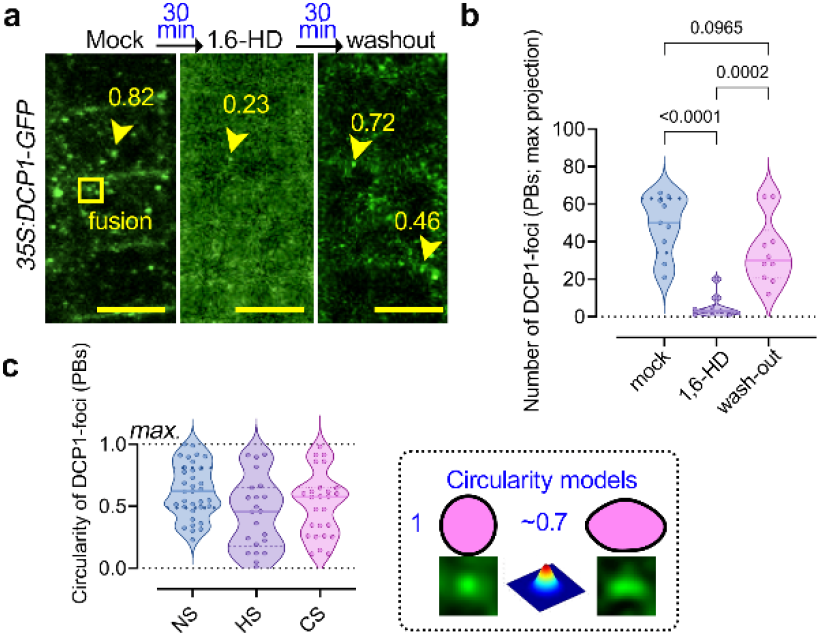
Effects of hexanediol, HS, CS and NS in circularity of PBs. **a** Treatment with 1,6-hexanediol induces the dissolution of PBs in the epidermal root meristematic region. Most PBs are sensitive to the aliphatic alcohol 1,6-hexanediol, which reversibly dissolves most LLPS-formed droplets^57^. After washout, DCP1-positive PBs reassemble, as indicated by representative confocal microscopy images; scale bars, 10 μm (*N*=2). Arrowheads denote circularity values for the indicated DCP1-positive PBs. Note that persistent PBs in the presence of 1,6-hexanediol show reduced circularity, suggesting that they likely are “hardened” through, e.g., Ostwald ripening. Hexanediol can compromise membrane integrity and lead to cytotoxic effects. Hence, the washout experiments can provide evidence for cell viability^58^. A fusion event is also shown (mock). We did not observe fusion/fission of PBs following 1,6-hexanediol addition, suggesting that the remaining PBs undergo coarsening. **b, c** Number and circularity of PBs decorated with DCP1-GFP (*35S:DCP1-GFP* transgene) in the presence/absence of 1,6-hexanediol (**b**) or at different temperatures (**c**) (*N*=1, *n*=11, ordinary one-way ANOVA). Circularity was determined as offsets from a Gaussian pixel spread function (above the graph; circularity = 4π x [area/perimeter^2^]). A circularity value of1.0 indicates a perfect circle. As circularity approaches 0.0, it indicates an increasingly elongated polygon. Values may not be valid for very small particles, which were excluded from our analyses (close to the diffraction limit of confocal microscopy, ∼200 nm). PB dissolution was also enhanced by cold stress (CS; 4°C), while PB formation increased upon heat stress (HS; 37°C, 2 h) (*N*=1, *n*=11, significant difference determined by ordinary one-way ANOVA). This observation suggests that DCP1 or many PB components, in general, show lower critical solution temperatures, which is consistent with its amino acid content (DCP1 is miscible below a certain temperature)^59^. Together with Supplementary Fig. 1, these results suggest that DCP1-positive PBs are LLPS condensates.

**Supplementary Fig. 3.**
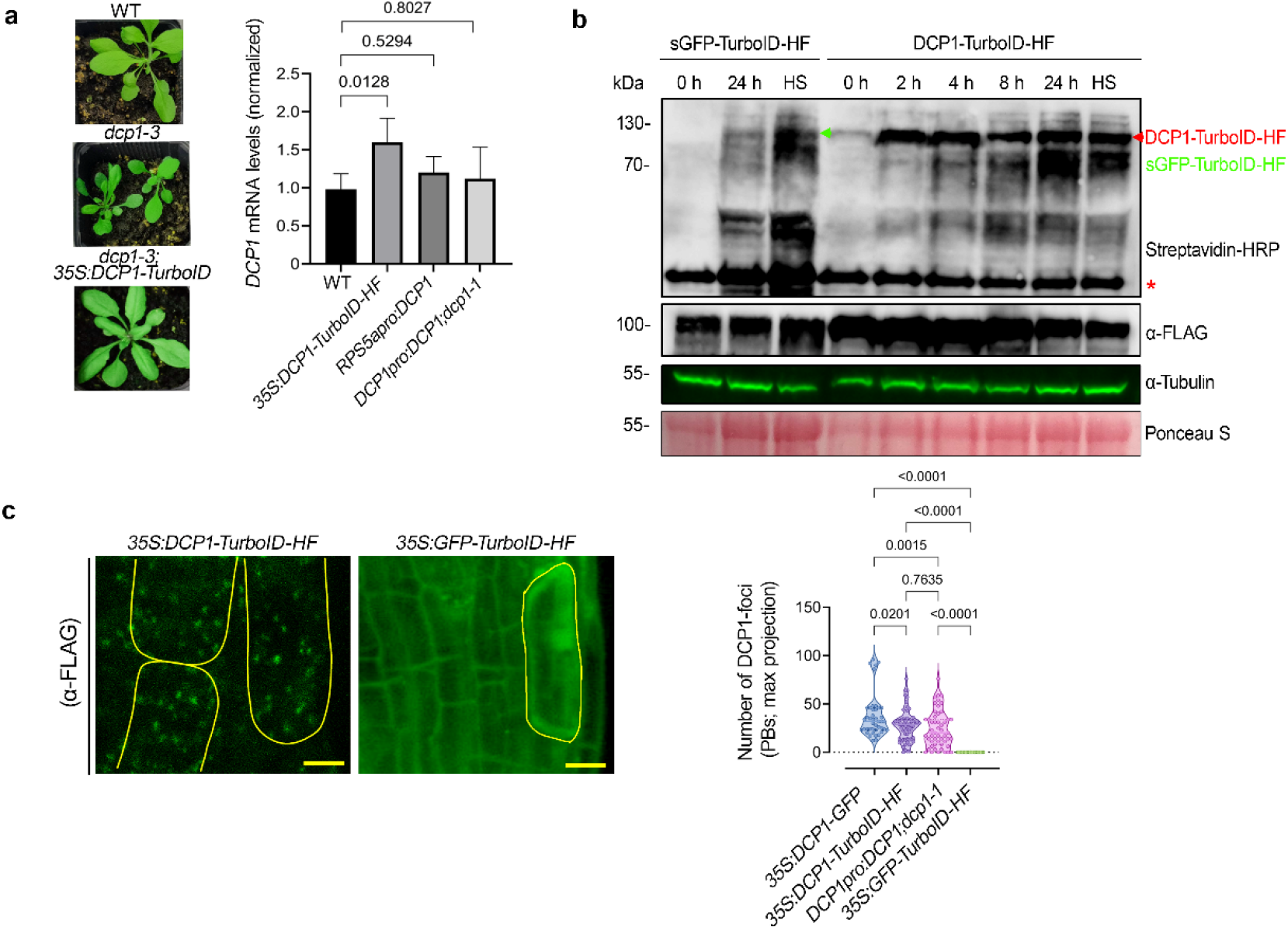
Establishment of a functional DCP1 bait for PDL. **a** The *DCP1-TurboID-HF* construct efficiently rescues the adult *dcp1-3* mutant phenotype. Left: phenotypes (3-week-old) of adult plants expressing *35S:DCP1-TurboID-HF* in the *dcp1-3* mutant background. In a semi-controlled greenhouse setting (temperature control 25°C), the *dcp1-3* mutant showed a smaller adult stature, as shown. Right: RT-qPCR analyses for the quantification of at *DCP1* expression levels (1-week-old whole seedlings). Data are means±SD (*N*=2, *n*=6; significant difference determined by *t-*test). **b** Representative confocal micrographs showing that the localization of DCP1 (anti-FLAG) to PBs is retained in lines expressing *DCP1-TurboID-HF*. As a control, the *GFP-TurboID-HF* line was used that does not show localization to PBs. Right: number of DCP1-positive foci in the corresponding lines expressing *DCP1*. Data are means±SD (*N*=1, *n*=32–60, significant difference determined by ordinary one-way ANOVA). **c** Immunoblot analyses of lines expressing *sGFP-TurboID-HF* or *DCP1-TurboID-HF* upon NS or HS conditions (same as used for APEAL). The immunoblots also show the accumulation of auto-biotinylated sGFP-TurboID-HF or DCP1-TurboID-HF in a time course of biotin administration (as detected with streptavidin-HRP, which captures biotinylated proteins). 50 µM Biotin was delivered in leaves by syringe infiltration and diffusion. Note that HS did not increase the biotinylation efficiency of DCP1-TurboID-HF: at 24 h, compare samples “2” with “4” in the “dynaIP”. HF, 6xhis-3xFLAG. The red arrowhead indicates the position of DCP1-TurboID-HF. Data are from a single experiment replicated twice. We used the same scheme for biotin application as determined in ref.^6^. The 2 h NS/HS corresponds to the timing after the administration of biotin for 24 h (*t*=0 corresponds to 24 h biotin administration in NS conditions). The red asterisks indicate non-specific bands.

**Supplementary Fig. 4.**
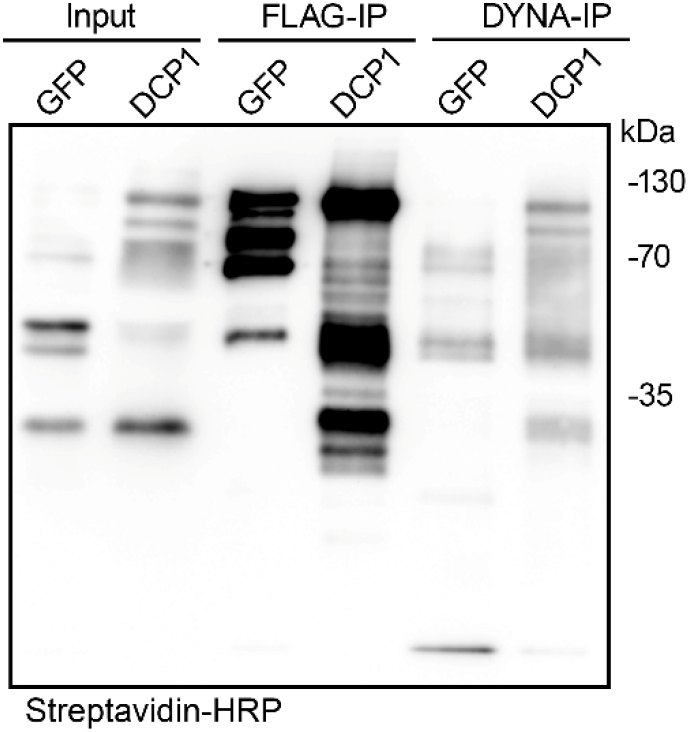
Biotinylated proteins evade purification from the AP step in APEAL. Immunoblot analyses from lines expressing *GFP-TurboID-HF* or *DCP1-TurboID-HF* (denoted as GFP or DCP1, respectively) showing the presence of biotinylated proteins in the flow-through after the AP-step (AP; for flow-through 1, see DYNA-IP here). The administration of biotin was performed as described in Supplementary Fig. 3 (no vacuum). The blot was overexposed to detect the faint streptavidin smear in flow-through 1. The baits in these experiments undergo auto-biotinylation, as reported previously. The results are representative of one experiment performed three times. Note that the immunodetectable biotinylation levels did not correlate well with the hits identified in Supplementary Fig. 5 (compare GFP to DCP1). GFP and DCP1 are of similar molecular weight, while the bands below the upper band (∼130 kDa) likely correspond to proteolytic cleavage products.

**Supplementary Fig. 5.**
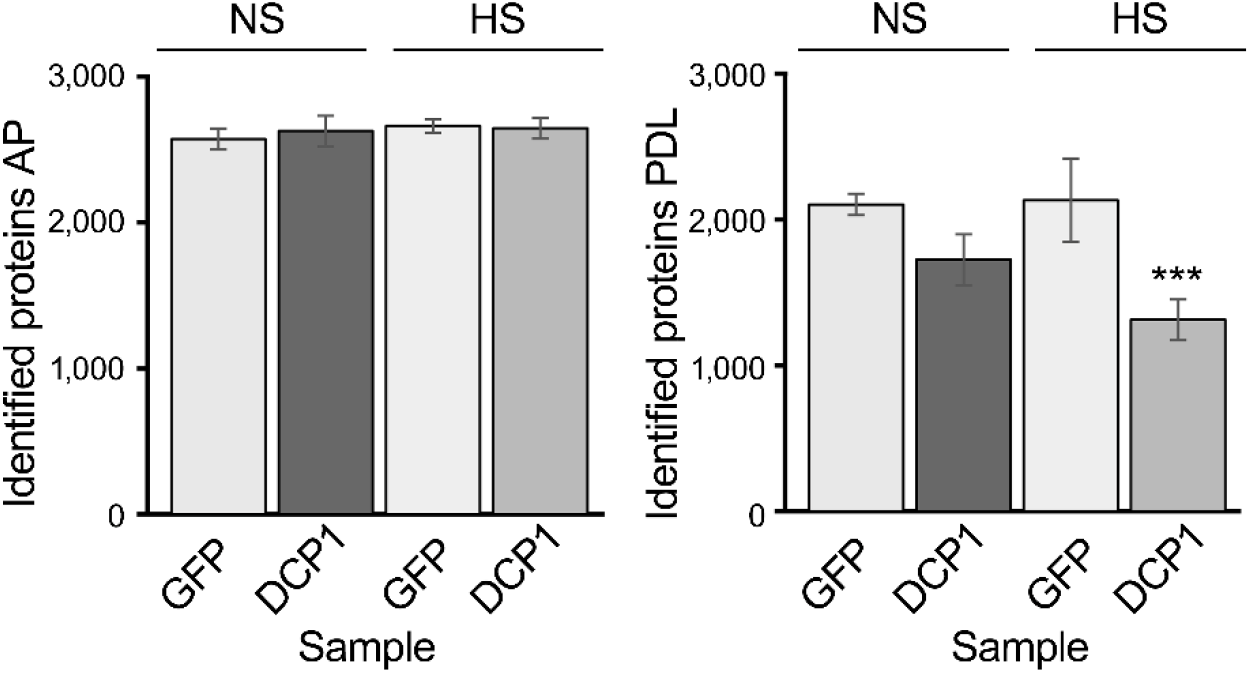
Hits from the APEAL approach (AP and PDL steps). Total protein hits from mass spectrometry under NS or HS conditions following the AP (left) or PDL (right) steps of APEAL. Proteins were extracted from lines expressing *sGFP-TurboID-HF* or *DCP1-TurboID-HF* (denoted as GFP or DCP1, respectively). The administration of biotin was as described in Supplementary Fig. 4; we used the same scheme for biotin application, as described in^6^: 24 h of 50 μM biotin, delivered without vacuum (as vacuum did not affect the overall profile of biotinylation; Supplementary Fig. 3). The 2-h NS/HS corresponds to the timing after the administration of biotin for 24 h (*t*=0 corresponds to 24 h biotin administration at NS). The results presented are unfiltered, containing the noisy portion of the proteome. Note that the free diffusion of GFP *in vivo* led to increased proteins identified. *sGFP-TurboID-HF*, GFP; *DCP1-TurboID-HF*, DCP1. Data are means±SD (*N*=3; \*\*\**p*≤0.05, as determined by *t*-test for comparison between HS and NS). Note the increased numbers of hits in GFP/PDL reflect the noisy proteome. GFP is expected to produce more noise (translated as hits in the context of the proteome), due to the increased diffusion over the specifically and topologically restricted DCP1 (confirmed in Supplementary Fig. 3, localizations). The decreased number of hits for HS conditions corresponds mainly to proteins of signal transduction and metabolism, as well as vesicle trafficking proteins (see also Fig. 2 and below for an explanation: HS reduces the DCP1 association with the PM). Furthermore, DCP1, as described below, loses localization at the PM during HS conditions. The AP/PDL GFP samples did not differ at *p*<0.005.

**Supplementary Fig. 6.**
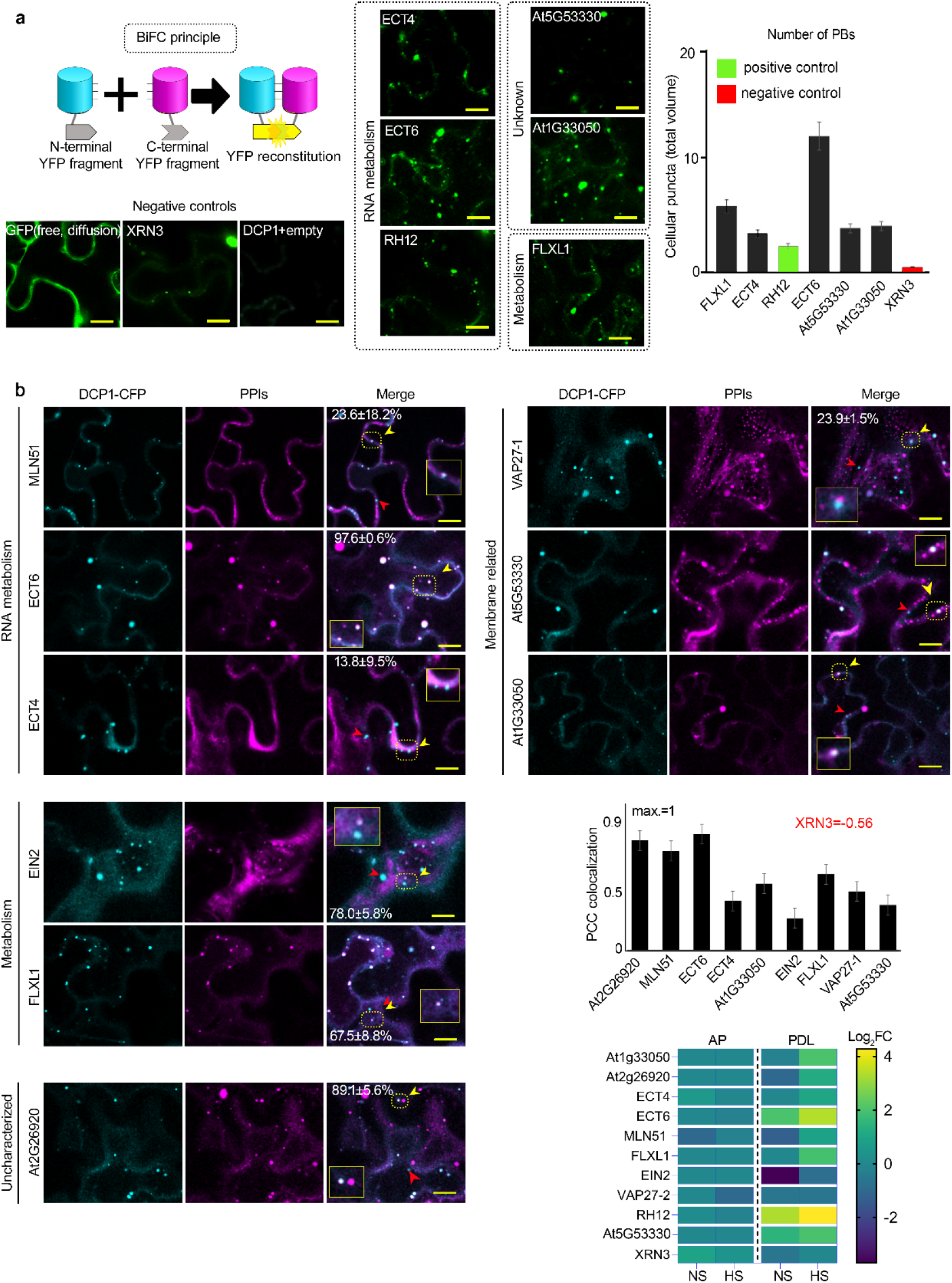
DCP1 colocalization and association with novel and known interactors in transient infiltration of *N. benthamiana* leaves. **a** Bimolecular fluorescence complementation (BiFC) assay of the indicated proteins. For protein selection, we classified the associations with DCP1, in both AP and PDL steps, according to their relative enrichment (log_2_FC). We selected proteins presenting moderate relative enrichment in the AP step, while having high predictability in the PDL step (Log_2_FC>0 in both steps). As an additional filter, we selected proteins rich in IDRs (half of the proteins from the list have high prion-like domain (PRD) scores and high Finum [aa]/total [aa] ratios; Supplementary File 5), and as such proteins would be likely to localize to condensates. We tested the association between PBs (using DCP1 as one representative component) and selected six proteins from these bins using BiFC and colocalization assays in *Nicotiana benthamiana* leaves transiently expressing pairs of appropriate constructs (Supplementary Fig. 6a,b; Supplementary File 5). These results suggested that the APEAL approach may help identify PB components or DCP1 interactors. These interactions should be further studied in Arabidopsis. BiFC efficiency was estimated from the reconstituted YFP raw signal intensity and YFP-positive puncta per cell in maximum projections (*N*=3). When two proteins interact, the cYFP and nYFP halves are brought in close proximity and produce a fluorescent signal. The diagram on the top depicts the BiFC concept and YFP reconstitution caused by protein-protein interaction *in vivo*. Lower left: representative confocal micrographs showing that YFP signal is reconstituted at cellular puncta that most likely correspond to PBs. XRN3 represents a negative control (see also **b**), as it localizes in the nucleus. Scale bars, 7 μm. Right: number of cellular puncta per total cell volume (in maximum projection images). Data are means±SD (*N*=3, *n*=20 cells). **b** Colocalization of selected proteins with DCP1-CFP-positive puncta (*35S:DCP1-CFP* transgene). The coding sequences of the corresponding “interactors” (PPIs; direct or indirect, defined in APEAL) were driven by the *35S* promoter and cloned in frame with *mCherry* at their 5′ end. Two-color colocalizations were estimated by Pearson’s correlation coefficients (PCC) using ultra-fast super-resolution microscopy combined with image deconvolution (∼120 nm axial resolution). Numbers in “merge” indicate colocalization frequency (mean)±SE between DCP1-CFP and the corresponding interacting protein (*N*=2, *n*=5 cells). Yellow arrowheads indicate colocalization and red arrowheads lack of colocalization. Lower right: PCC of pixel intensities between DCP1-CFP and the corresponding putative interacting protein. Data are means±SD (*N*=3, *n*=20 cells). We confirmed the ECT domain-containing proteins, MLN51, FLXL1, EIN2, VAP27-1 and uncharacterized At1g33050 (hypothetical protein) and At5g53330/At2g26020 (ubiquitin-associated/translation elongation factors EF1B) as novel PB components. By applying a pre-selection criterion of enrichment (log_2_FC>0.5) for the selection of preys, we significantly increased the probability of identifying successful binary interactions between PBs (i.e., cYFP-DCP1) and the identified proteins. In particular, all preys were confirmed as PB components (although FLXL1 is marginal), while PDL had higher interaction predictive power than the AP step irrespective of whether proteins were enriched in NS or HS (Supplementary File 5). XRN3 was used as a threshold control (log_2_FC=–0.58, PDL/NS conditions). The heatmap shows the enrichment of these proteins in the different conditions; the scale at right shows log_2_FC.

**Supplementary Fig. 7.**
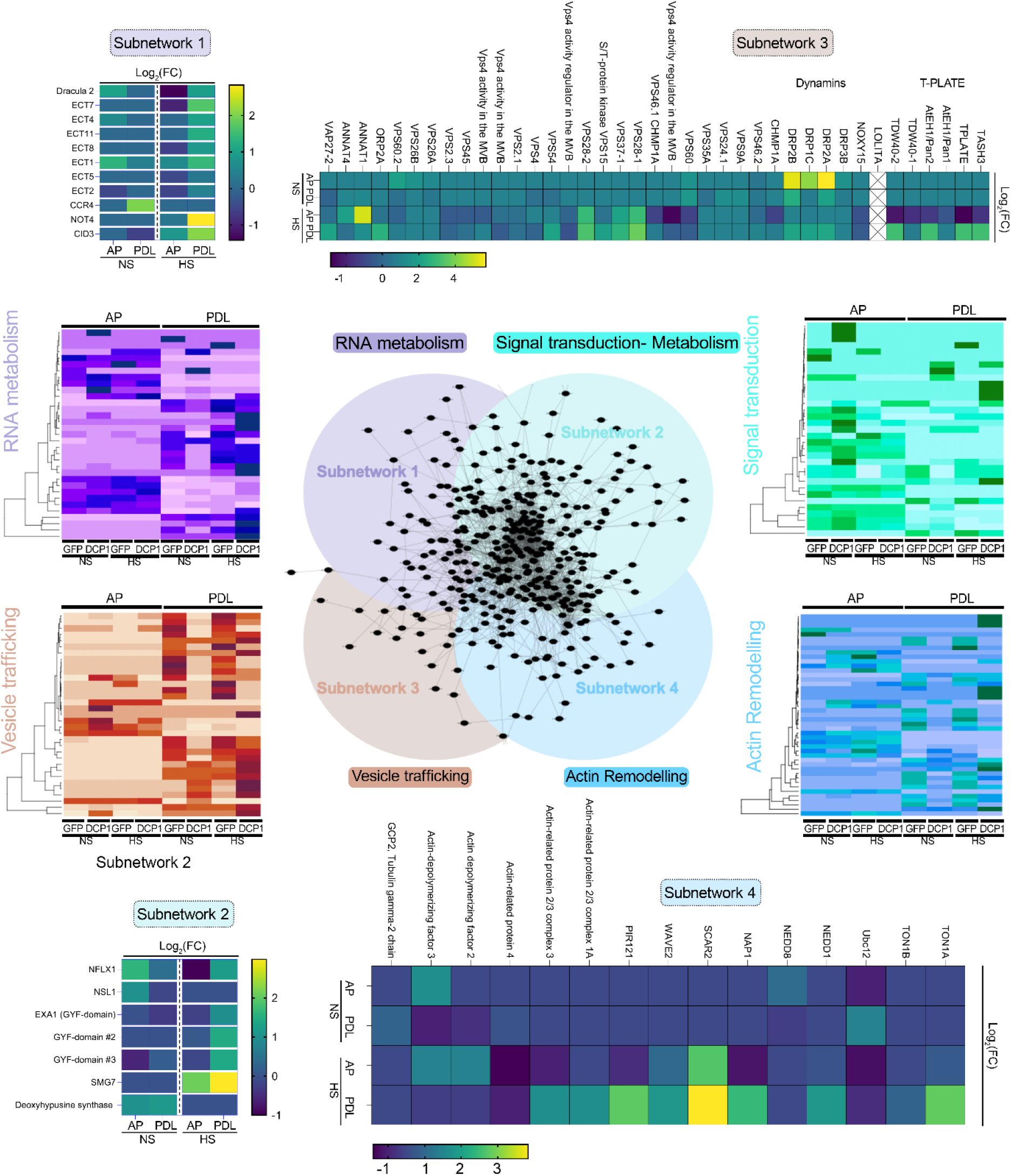
Structure and density of the four interconnected networks. Subcluster analyses of APEAL reveal four interconnected subnetworks (center). The four networks were: proteins related to RNA metabolism; signal attenuation and metabolism; vesicle trafficking; and actin remodeling. Heatmaps depict the abundance of selected proteins from the four subnetworks. Notably, the AP step in the “purple” heatmap did not lead to the identification of RNA metabolism proteins. Below, we summarize interesting hits from each network: **Subnetwork 1: RNA metabolism** Nuclear pore complex (NPC) proteins engage in processing, capping, splicing and polyadenylation; NPC proteins were enriched in PDL/HS. For example, DRACULA (DRA2) is a homolog of mammalian NUP98 and a likely NPC component in Arabidopsis ^60^. In *Caenorhabditis elegans*, perinuclear P granules contain components of PBs and SGs; P granules at NPC clusters might interact with nascent RNAs^61^. As the interaction of PBs with NPC occurs under HS conditions, PBs may be recruited there to store nascent RNAs or block them from nuclear export. Furthermore, mRNAs undergo several modifications; one of the most prevalent is *N*^6^-methyladenosine (m^6^A), which is of key importance in development. Both AP/PDL datasets (mainly HS; Supplementary File 6) contained the YTH (YT521-B homology)- domain ECT (EVOLUTIONARILY CONSERVED C-TERMINAL REGION) proteins, which are praise *bona fide* m^6^A readers controlling leaf development^62^. The ECT animal homologs, YTHDF1/2/3, undergo LLPS induced by polymethylated mRNAs acting as a multivalent scaffold for their binding, juxtaposing their low-complexity domains and thereby leading to LLPS^63^. We identified components of the adenylation regulatory machinery, including the CARBON CATABOLITE REPRESSION 4 (CCR4)–NOT complex and POLYADENYLATE BINDING PROTEINs (PABs). For PDL/HS, we identified the PAB-interacting protein CTC-INTERACTING DOMAIN 3 (CID3), a homolog of ataxin-2 identified in mammalian PBs^64^. At3g45630 is a putative ubiquitin (Ub) E3 ligase with an RRM (RNA-recognition motif)^65^, showing architectural similarity to the RING E3 ligase NOT4; we confirmed through BLAST-P analysis its orthology to human NOT4 (*p*=6e^−59^), which catalyzes the removal of the poly(A) tail. Likewise, CCR4 localizes to plant PBs^66^. Ub, as well as the proteasome itself, enters condensates^67^, and thus it is not surprising that E3 ligases are recruited to PBs. Furthermore, we identified argonaute1 (AGO1) in the PDL (Fig. 1b), involved in RNA metabolism. As AGO1 is involved in nascent peptide ubiquitination^68^, this function could also involve PBs and associated E3 ligases. **Subnetwork 2: Defense Regulation and Metabolism** The NF-X1-type zinc finger protein NFXL1 (PDL/HS) functions as a negative regulator of defense-related genes via a salicylate (SA)-dependent signaling pathway. Likewise, NECROTIC SPOTTED LESIONS 1 (NSL1), a MACPF domain-containing protein, negatively controls SA-mediated cell death^69^. We also identified three proteins with glycine-tyrosine-phenylalanine (GYF; PDL/HS) domains; one, Essential for poteXvirus Accumulation 1 (EXA1), recognizes Pro-rich sequences, as do other GYF domain-containing proteins found in humans and yeast^70^. EXA1 is indispensable for *Plantago asiatica* mosaic virus infection and negatively regulates levels of plant immune receptors via translational repression. EXA1 also interacts with SMG7 (also enriched in the PDL/HS), which plays a role in nonsense-mediated RNA decay (NMD)^70^. Two additional GYF-containing proteins identified are of unknown function (At1g24300 and At1g27430). These findings suggest that PBs restrain cell death via immune signaling and may function as a hub in SA perception. We identified proteins involved in translational regulation, e.g., deoxyhypusine synthase (At5g05920; AP)^71^. This enzyme mediates the post-translational activation of eIF5A, which facilitates the translation of specific mRNAs. This confinement in PBs may partition deoxyhypusine synthase into an inactive pool to reduce translation and minimize energy squandering during stress. **Subnetwork 3: Membrane remodeling/trafficking** Unexpectedly, we detected significant enrichment for membrane-associated proteins in the PDL proxitome. This subnetwork was further divided into four major dense clades consisting of 1) ESCRT (endosomal sorting complex required for transport), 2) dynamin-related proteins (DRPs; see also Fig. 2a) and 3) the plant-specific endocytic TPLATE complex. DRPs are force-generating proteins targeted to membranes that regulate membrane remodeling pathways^72^. We observed an enrichment of DRPs (i.e., 2A/B/E/C/E, ARC5 and RSW9) for both AP and PDL. Furthermore, DRPs are physically linked to the TPLATE complex^6^, and enriched under HS conditions (e.g., AtEH2, log_2_FC_NS/HS_=0/3.1). Recently AtEH1 was shown to phase-separate to induce endocytosis^73^. We hypothesize that during stress, DCP1 may cooperate with the endocytotic machinery. This hypothesis is possible, considering that eisosomes, static structures that arguably modulate endocytosis, contain the XRN4 yeast homolog XRN1^74^. The complexes ESCRT-0 -I, -II, and -III drive Ub-mediated cargo sorting from the PM^75^. ESCRT components were enriched in the PDL: e.g., ELC (Vps23p/TSG101, multivesicular bodies [MVB] sorting), VPS28/37 (ESCRTI complex) containing a non-canonical RNA-binding motif, and the VPS15, which sort endocytic cargos into MVBs. In animals, ESCRT components have been found in condensates^76^. DCP1 or the PBs may aid in the formation of MVBs or load them with specific RNA molecules. Furthermore, we identified proteins residing in interorganellar contact sites: for example, Annexin 1 (ANNAT1; PDL/NS; 17 peptides vs. 0 in the control). In animals, Annexin A11 (ANXA11) is an RNA granule-associated phosphoinositide-binding protein that functions as a molecular tether between RNPs and membranes^77^. Likewise, in the endosome-vesicular trafficking proteins, the ORP2A, OSBP (OXYSTEROL BINDING PROTEIN)-RELATED PROTEIN ORP-related Osh lipid exchange proteins were found enriched in PDL. ORPs create a potential nanoscale membrane lipid environment^78^ controlling PM organization and dynamics and marking ER-PM contact sites (ERPC), while in animal cells LLPS in ORP-decorated sites marks contacts with peroxisomes and lysosomes^37, 79^. **Subnetwork 4: Novel Interactors: Actin polymerization and cytoskeleton remodeling** With regards to actin, we identified SCAR–WAVE components (cluster score *p*=5.41e^−5^; FDR=0.05) and accessory regulators like PIR121 and NAP125 (NAP1; both in PDL/HS, see also below), which collectively modulate actin/microtubules dynamics^80^. Likewise, for microtubules (MTs), we identified nucleators like TONNEAU1 (TON1) that share similarities with the human centrosomal protein FOP (FGFR1 Oncogene Partner). TON1 interacts with a superfamily of 34 proteins conserved in land plants that harbor the TON1A/B Recruiting Motif (TRM) involved in spindle and phragmoplast organization^81^. TRM proteins involve the evolutionarily conserved γ-tubulin complex (TuRC); in agreement with this, we identified NEDD1, which interacts with TuRC ^82^. In animals and fungi, γ-tubulin is a nucleator localizing in the MT organizing center (the centrosome in animal cells and the spindle pole body in yeast).

**Supplementary Fig. 8.**
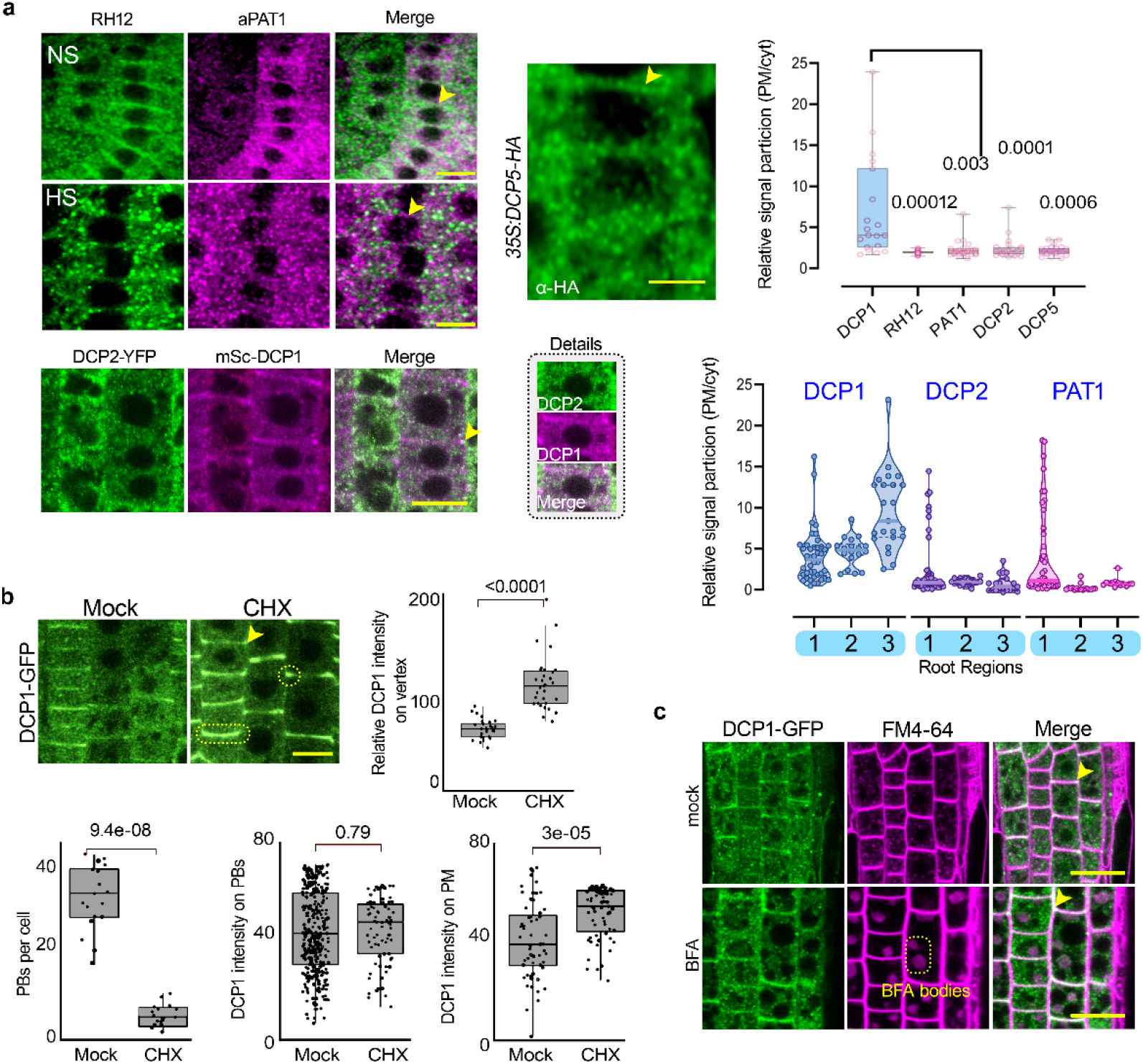
Core PB components tend to localize at the PM and DCP1 localization upon dissolution of PBs. **a** Representative confocal micrographs showing the transient PM localization of DCP1, RH12, PAT1 (detected by anti-PAT1), DCP2 and DCP5 under NS or HS conditions (for RH12 and PAT1). The micrographs correspond to region 1 of epidermal cells. Note the absence of DCP1/DCP2 colocalization at the PM (or edge/vertex, as discussed below) in lines coexpressing *RPS5apro:mScarlet-DCP1* and *35S:DCP2-YFP* (epidermis cells, region 2). A detailed inset on the right is also shown. The arrowhead points to the likely localization of DCP1 at the PM. Scale bars, 5 μm. Right: box plot showing the PM-to-cytoplasmic signal ratio (upper). The ratio of the signal here is approximate, as the PM was defined as a line along the cell contours. Data are means±SD (*N*=3, *n*=13–42 cells, significant difference determined by nested one-way ANOVA). Bottom right: PM-to-cytoplasmic signal ratio in three root regions of DCP1 (from the *DCP1pro:DCP1-GFP* transgene), DCP2 (*35S:DCP2-YFP*) and PAT1 (*PAT1pro:PAT1-GFP*) (*N*=3, *n*=13–42 cells). **b** Representative confocal micrographs showing that cycloheximide (CHX) treatment does not result in the release of DCP1-GFP from the PM, edges/vertices (region 2, epidermis). Scale bar, 20 μm. The boxplots show DCP1-GFP intensity on edge/vertex, PM, PBs, and DCP1-positive PBs number under control or CHX conditions (50 μm, 30 min). Data are means±SD (*N*=3, *n*=33 cells, Wilcoxon). DCP1 intensity in PBs remaining after CHX was calculated in DCP1-positive foci/remnants. **c** Representative confocal micrographs showing the lack of effect from brefeldin A (BFA; 50 μm for 1 h) treatment on the DCP1 route to the PM. Scale bars, 10 μm. Arrowhead (top) denotes the edge/vertex, and BFA bodies (stained by FM4-64) are at the bottom. BFA treatment in Arabidopsis roots causes the formation of TGN/endosomal agglomerates called BFA-induced compartments^31^.

**Supplementary Fig. 9.**
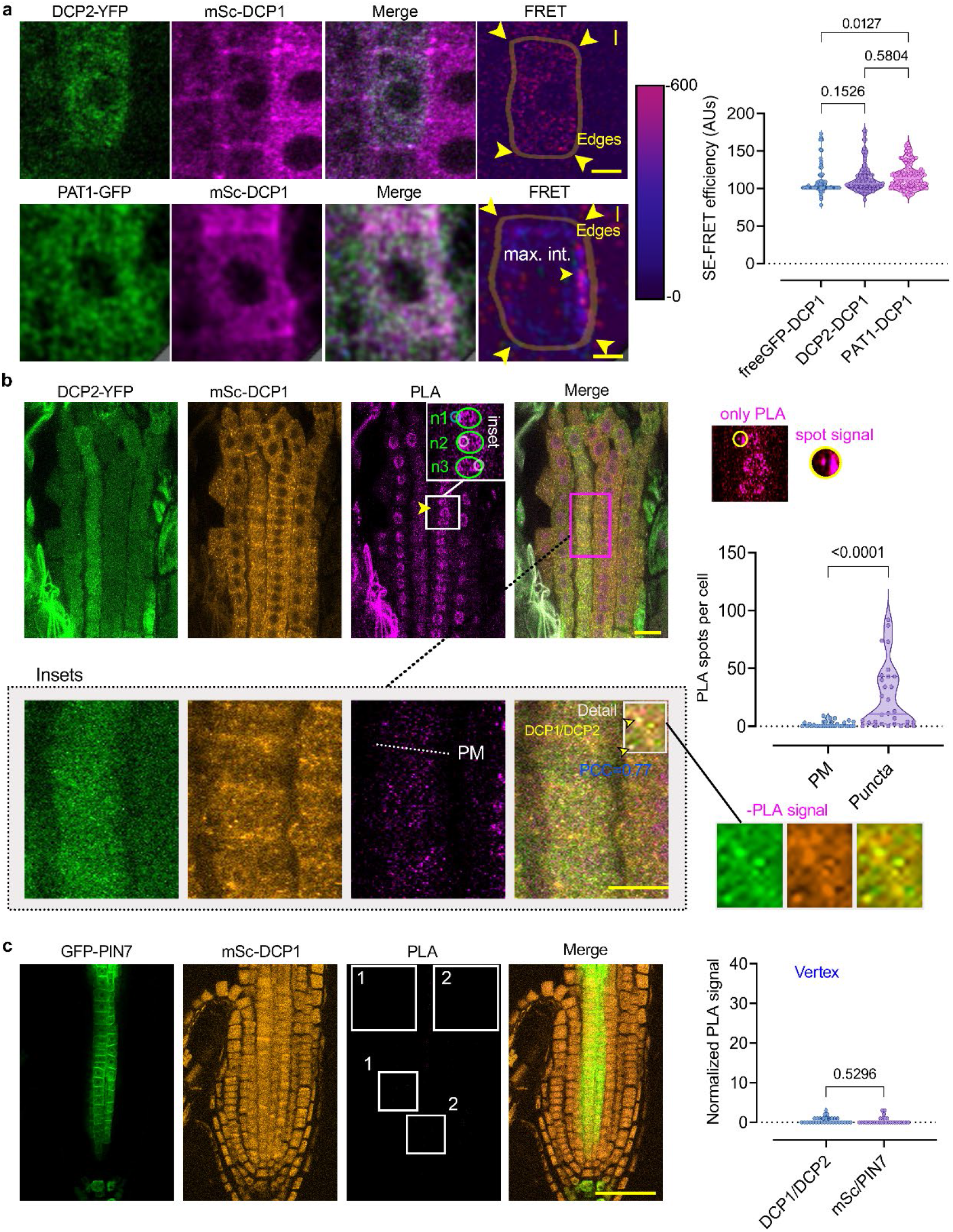
DCP1/DCP2 or DCP1/PAT1 spatiotemporal interaction by PLA and SE-FRET. **a** Sensitized emission (SE)-FRET efficiency between DCP2-YFP and mScarlet-DCP1 in lines co-expressing *RPS5apro:HF-mScarlet-DCP1* and *35S:DCP2-YFP* or *PAT1pro:PAT1-GFP* (cell vertices/edges; region 2, epidermal cells). Scale bars, 5 μm. The cell contours are shown. Right: signal quantification of SE-FRET efficiency between the indicated combinations (DCP2/DCP1 and DCP1/PAT1). As a negative control, lines co-expressing *GFP* and *DCP1* were used. Data are means ±SD (*N*=3, *n*=6–8, significant difference determined by Kruskal-Wallis). Note also that FRET was observed for puncta in the cytoplasm. Max. int., maximum FRET intensity observed (scale on the right). **b** Representative confocal micrographs and a graph showing PLA-positive signal (as spots per cell), produced by anti-FLAG/anti-GFP for the *HF-mScarlet-DCP1* + *DCP2-YFP* combination at the PM (all root regions including edge and vertex) and in the cytoplasm (puncta), respectively. The “n1–3” from the PLA panel, denotes three nuclei with positive PLA spots (inset in PLA; small circles denote the PLA spots). On the right, the inset is presented without labeling (only “PLA”), along with a spot detail. Note also the absence of PLA spots along the PM plane (shown as an inset below the main micrograph). PCC between DCP1 and DCP2 is also shown for the “detail”. Scale bars, 5 μm. **c** Representative confocal micrographs and a graph showing PLA-positive signal (as spots per cell), produced by anti-FLAG/anti-GFP for *the HF-mScarlet-DCP1* + *PIN7-GFP* combination. This combination represents an additional negative control. PIN7 is a PM protein lacking from the APEAL dataset. The insets (1, 2), show the lack of PLA spots. Some background noise is evident. Scale bar, 100 μm. Data are means ± SD (*N*=5, *n*=111, edges/vertices, significant difference determined by ordinary one-way ANOVA).

**Supplementary Fig. 10.**
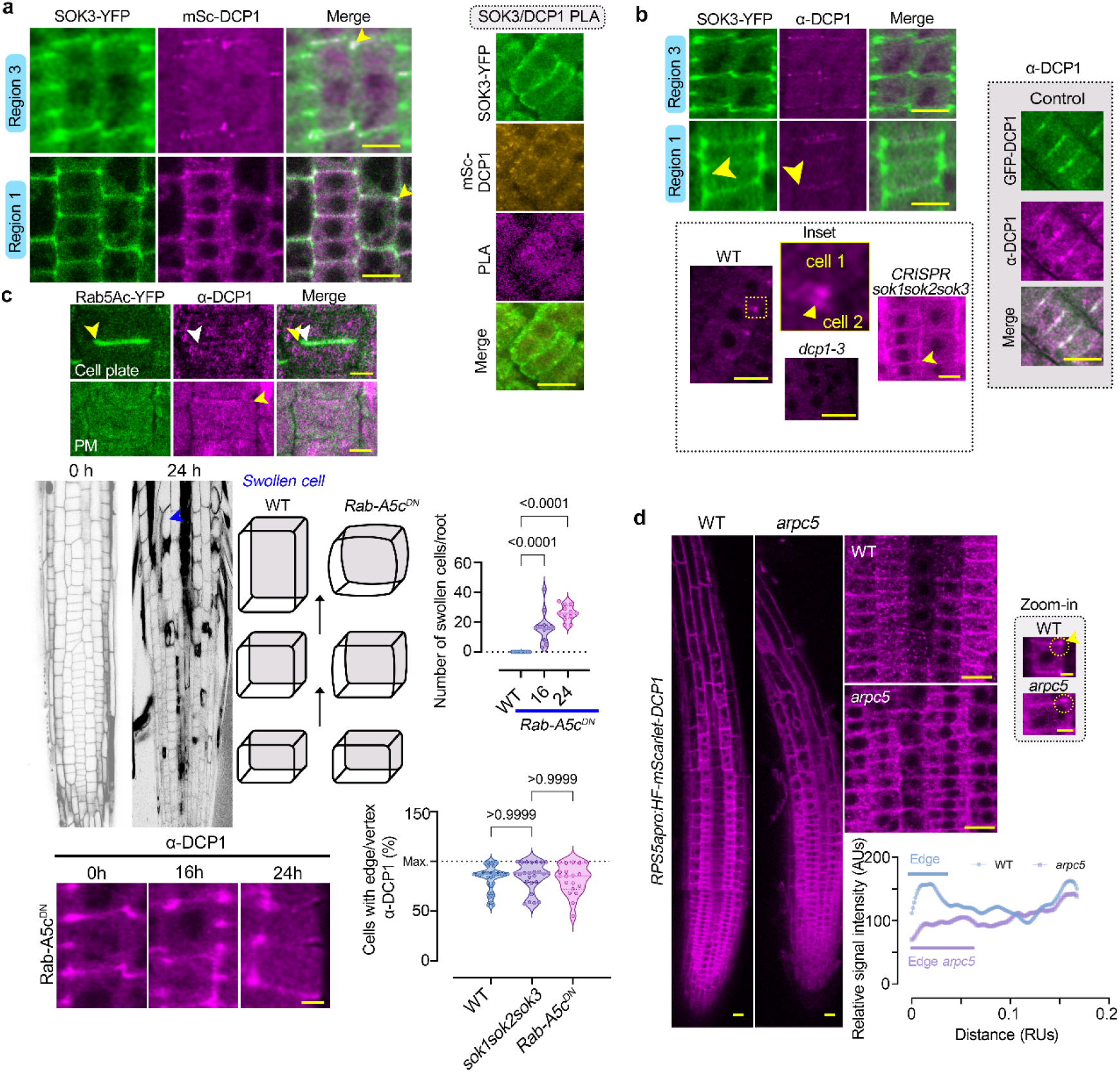
DCP1 is likely not recruited to the cell edge by Rab-A5c or SOK3. **a** Representative confocal micrographs showing the localization of DCP1-mScarlet (from the *RPS5apro:DCP1-mScarlet* transgene) and SOK3-YFP (*SOK3pro:SOK3-YFP* transgene) from two regions as indicated (SOK3; left, no abnormal phenotype was observed for the root even for lines expressing *SOK3pro:SOK3-YFP*, unlike what has been suggested for SOK1). The experiment was performed three times with similar results. Scale bars, 7 μm. Right: PLA showing a lack of interaction between DCP1 and SOK3 (*HF-mScarlet-DCP1* + *SOK3-YFP*; anti-FLAG + anti-GFP, *N*=5). This result suggests that PLA has a shorter interaction range than PDL. Scale bar, 10 μm. **b** Representative confocal micrographs showing the localization of DCP1, as detected by immunostaining with anti-DCP1 and SOK3-YFP (*SOK3pro:SOK3-YFP* transgene; left, no phenotype was observed for the root) from two regions corresponding to 1 and 2–3. Bottom: representative confocal micrograph showing that the loss of SOK3 function does not affect DCP1 localization. Scale bars, 7 μm. The inset shows a detail of the vertex localization of DCP1; the yellow arrowheads point to the gap between two vertices in adjacent cells. Scale bars, 10 μm. Right: Confocal micrographs showing the detection of DCP1 using an anti-DCP1 antibody in root meristematic epidermal cells of WT seedlings, *dcp1-3* seedlings (with reduced signal), *sok1sok2sok3 CRISPR*, or seedlings expressing *35S:GFP-DCP1* (positive control). **c** DCP1 and Rab5Ac do not colocalize in dividing cells. Representative confocal micrographs showing the localization of DCP1 (detected by anti-DCP1) and Rab5Ac-YFP in cytokinetic or mesophase cells (*N*=2). The white arrowhead denotes the accumulation of Rab5Ac-YFP at the leading edge. Note the lack of DCP1 localization at the leading edge (offset of the yellow arrowhead in “Merge”). In the micrographs below, the edge/vertex is denoted by an arrowhead. Scale bars, 2 μm. Middle: PI-stained roots showing the radial swelling of a whole root expressing the dexamethasone-inducible line *Dex>>RAB-A5c^DN^*. Under the same conditions, WT roots did not show any swelling. The diagram shows the progressive (an)isotropic growth between WT and *Rab-A5c* inducible lines, respectively. Next to the diagram, a quantification of swollen cells in lines expressing *RAB-A5c^DN^* is shown (*N*=2, *n*=10). Bottom: DCP1 detection with the anti-DCP1 antibody (raised against full-length DCP1 protein purified from *Escherichia coli*) shows no differential localization of the protein upon dexamethasone induction (2 μM for up to 48 h). Scale bars, 5 µm. The graph indicates the percentage of cells with edge/vertex localization of DCP1 (*N*=3, *n*=5, significant difference determined by *t*-test). **d** Representative confocal micrographs showing HF-mScarlet-DCP1 localization in WT and the *arpc5* mutant background. Scale bars, 10 μm. Vertex localization of mScarlet-DCP1 (whose encoding construct is driven by the meristem-specific *RPS5a* promoter) is not significantly reduced by the lack of the Arp2–Arp3 complex, yet the signal is less confined, suggesting that Arp2–Arp3 is required for DCP1 domain restriction at the vertex/edge. Here, the *arpc5* allele *crooked* was used. Scale bars, 3 μm.

**Supplementary Fig. 11.**
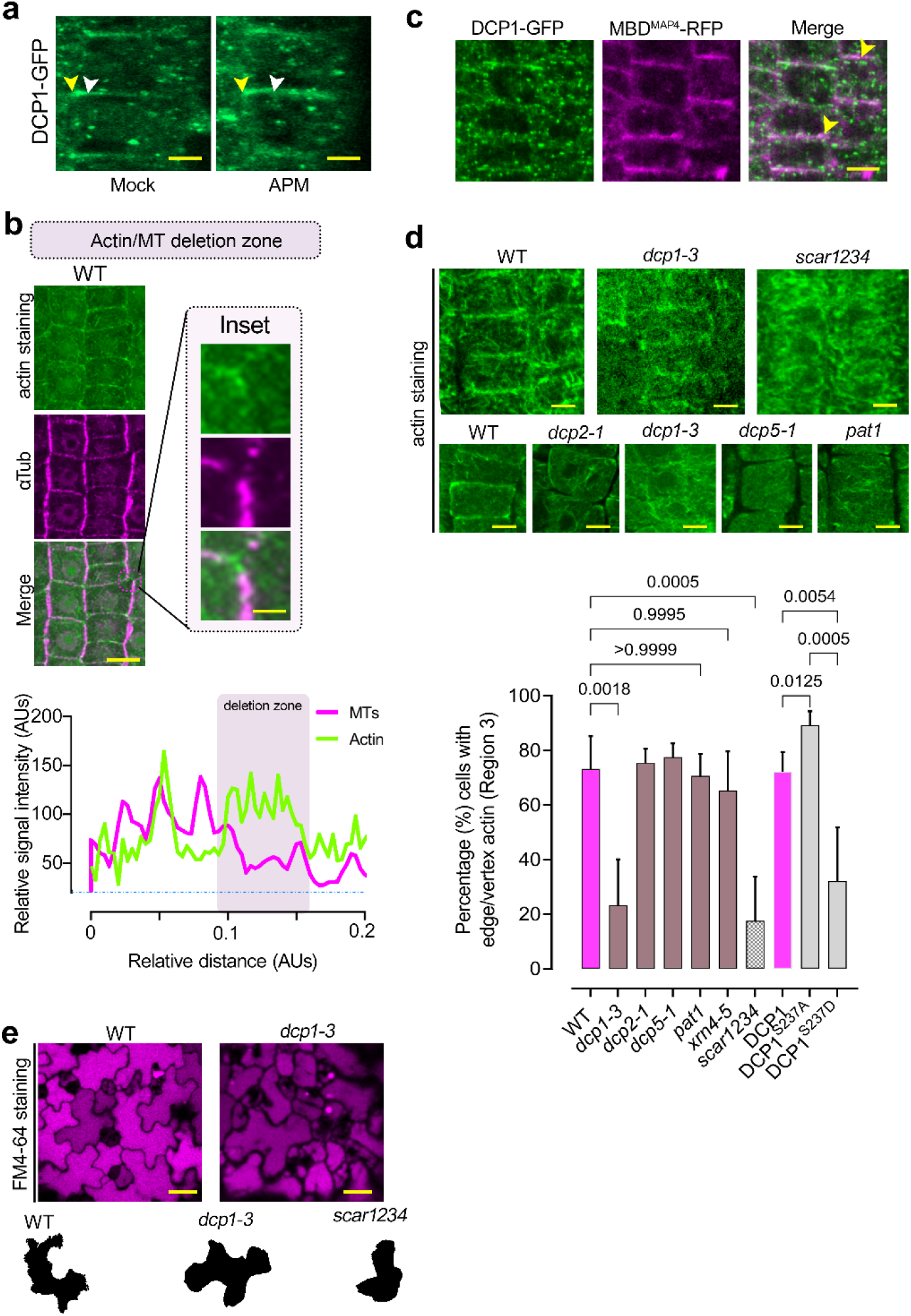
Actin localization in different mutants. **a** Effect of APM on DCP1 localization at edges/vertices. Representative confocal image showing the expansion of the DCP1 domain at edges/vertices (the white arrow indicates the expanding edge; epidermal cells of region 2). Scale bars, 5 μm. The expansion was 2.8±1.6 μm (*N*=5, *n*=10; region 2 epidermal cells). **b** Confocal micrographs showing root meristematic epidermal cells counterstained simultaneously with phalloidin (which marks actin) and with α-tubulin. Inset shows details of the edge. Note the cell vertex exclusion zone, where MTs are absent. Below: plot profile indicating the relative signal intensity of actin and MT staining along the cell edge. Scale bars, 10 μm. The MT deletion zone enriched in actin at the cell edges, as described previously for the trichome apex,^35^. This finding was important, as the loss of Arp2–Arp3 function leads to more variable MT deletion zones in other contexts^83^, and may explain the variable DCP1 domain size at edges/vertices in the absence of Arp2–Arp3 (Supplementary Fig. 10d). **c** Representative confocal micrographs showing DCP1/MAP-RFP exclusion from lines co-expressing *DCP1pro:DCP1-GFP* and the *MAP4^MBD^-RFP* microtubule marker line. Scale bar, 10 μm. **d** Representative confocal micrographs showing actin localization in WT, *dcp1-3*, *scar1234*, *dcp2-1*, and *pat1-1* upon phalloidin-actin staining. Scale bars, 10 µm. Bottom: percentage of cells in region 3 with accumulation of actin at edges in various genotypes. Scale bars, 5 μm. Data are means±SD (*N*=3, *n*=7–9 roots; significance was tested by Brown-Forsythe and Welch ANOVA; all 45 comparisons were determined, and the most important are shown). **e** Counterstaining of leaf epidermal cells with FM4-64 in WT and the *dcp1-1* mutant. Note the disruption of the jigsaw puzzle phenotype on the leaf surface, which resembles that described for *scar1234* or *arp2/3* complex mutants.

**Supplementary Fig. 12.**
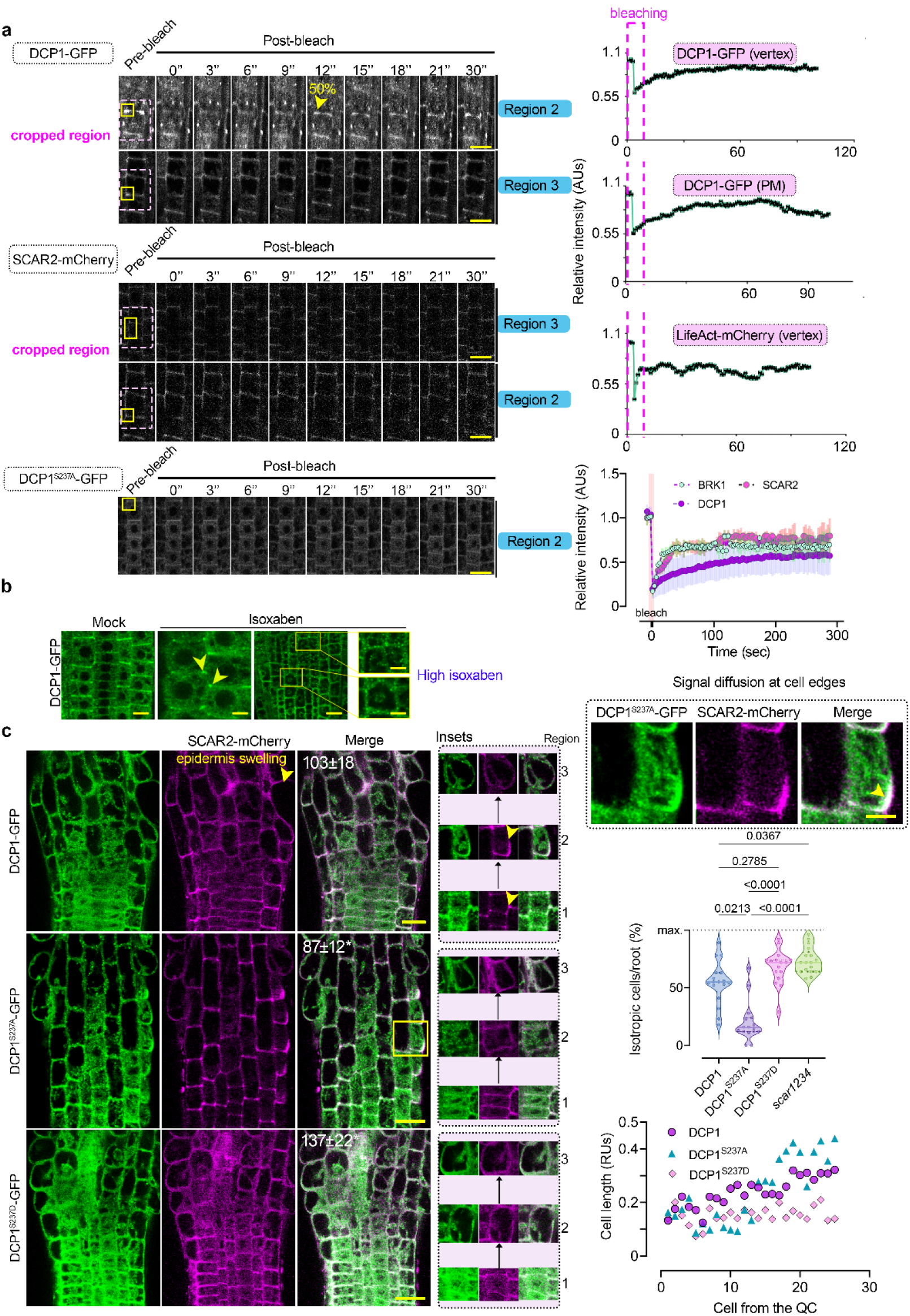
FRAP analyses of DCP1 in the edge/vertex and effect of isoxaben treatment on radial swelling (isotropic expansion) of phosphovariant DCP1 lines. **a** Representative confocal micrographs showing FRAP assays from regions 2 and 3. The cropped region used in Fig. 6 is indicated. Scale bars, 20 μm. The snapshot of 50% signal recovery is also shown (arrowhead in region 2; region 3 did not reach 50% of the initial signal in the given time). Right: recovery rates of DCP1-GFP at the PM and vertex, and of LifeAct-mCherry (vertex). Data in the lowest graph are means±SD (*N*=3, *n*=3 cells). AUs, arbitrary units. **b** Isoxaben (IXB, 30 μM, 8 h) slightly affected the vertex localization of DCP1 in some cells, PBs increased. Arrowheads denote vertex localization of DCP1. Epidermal cells, region 3; Scale bars, 10 μm/5 μm. **c** Representative confocal micrographs showing DCP1-GFP localization (and phosphovariants) in the corresponding root meristematic cells treated with isoxaben (IXB, 30 μM, 8 h). Insets indicate treated root cells along the different regions. Note the diffusion of SCAR2 and DCP1-GFP in cells that due to IXB treatment transition from anisotropic to isotropic expansion (arrowhead in the inset “signal diffusion at the cell edges”). The numbers in merge are means±SD (*N*=3, *n*=9 roots; asterisks indicate significance at *p*<0.05 significant difference determined by *t*-test) of root width. Scale bars, 20 μm. Furthermore, note the detail of DCP1/SCAR2 colocalization (denoted by the arrowhead). Lower right: percentage of cells showing isotropic growth and their distance from the quiescent center (QC) compared to their length, upon drug treatment (*N*=2, *n*=11–21, significant difference determined by ordinary one-way ANOVA).

